# Scalable hardware and software for high-density, multi-regional electrophysiology in non-human primates

**DOI:** 10.64898/2026.10.22.761135

**Authors:** Agrita Dubey, Qiao Xu, Mohammad Khazali, John Choi, Katie Wingel, Jarl Haggerty, Jacob Stefanowicz, Bijan Pesaran

## Abstract

Understanding how distributed neural circuits generate complex behaviors requires long-duration recordings from populations of individual neurons across multiple brain regions. High-density Neuropixels probes have revolutionized rodent systems neuroscience. Extending these capabilities to non-human primates (NHPs) for chronic, multi-regional recordings requires compact, implantable microdrive hardware and reliable automated spike-sorting software. NHPs provide access to complex and flexible behaviors that are difficult to study in other model systems. Extending the Neuropixels revolution to NHPs therefore depends on integrating hardware and software for long-duration, multi-regional recordings.

Here, we present a scalable hardware–software platform co-designed for high-density neural data acquisition and reliable single-unit identification in NHPs. The Multi-regional Implantable NeuroDrive (MIND) is a lightweight, implantable microdrive that provides stable recordings from multiple independently positioned Neuropixels probes and is designed to support chronic, ambulatory, head-free recordings. We paired MIND with UnitAssay, an automated post-sorting curation framework that identifies and curates characteristic spike-sorting errors with minimal computational cost. Together, MIND and UnitAssay provide an integrated platform for implanted, multi-regional recordings in NHPs performing complex and naturalistic behaviors.

We tested MIND and UnitAssay across parietal, prefrontal, and motor cortices in rhesus macaques performing reaching and eye-movement tasks. MIND enabled stable multi-probe recordings across cortical regions and durations. UnitAssay enabled automated curation of large-scale spike-sorting output and improved reliable single-unit identification relative to Kilosort 4 labels and standard SpikeInterface quality control. Together, MIND and UnitAssay provide a scalable solution for high-density, multi-regional, long-duration electrophysiology in NHPs.

**Graphical Abstract:** 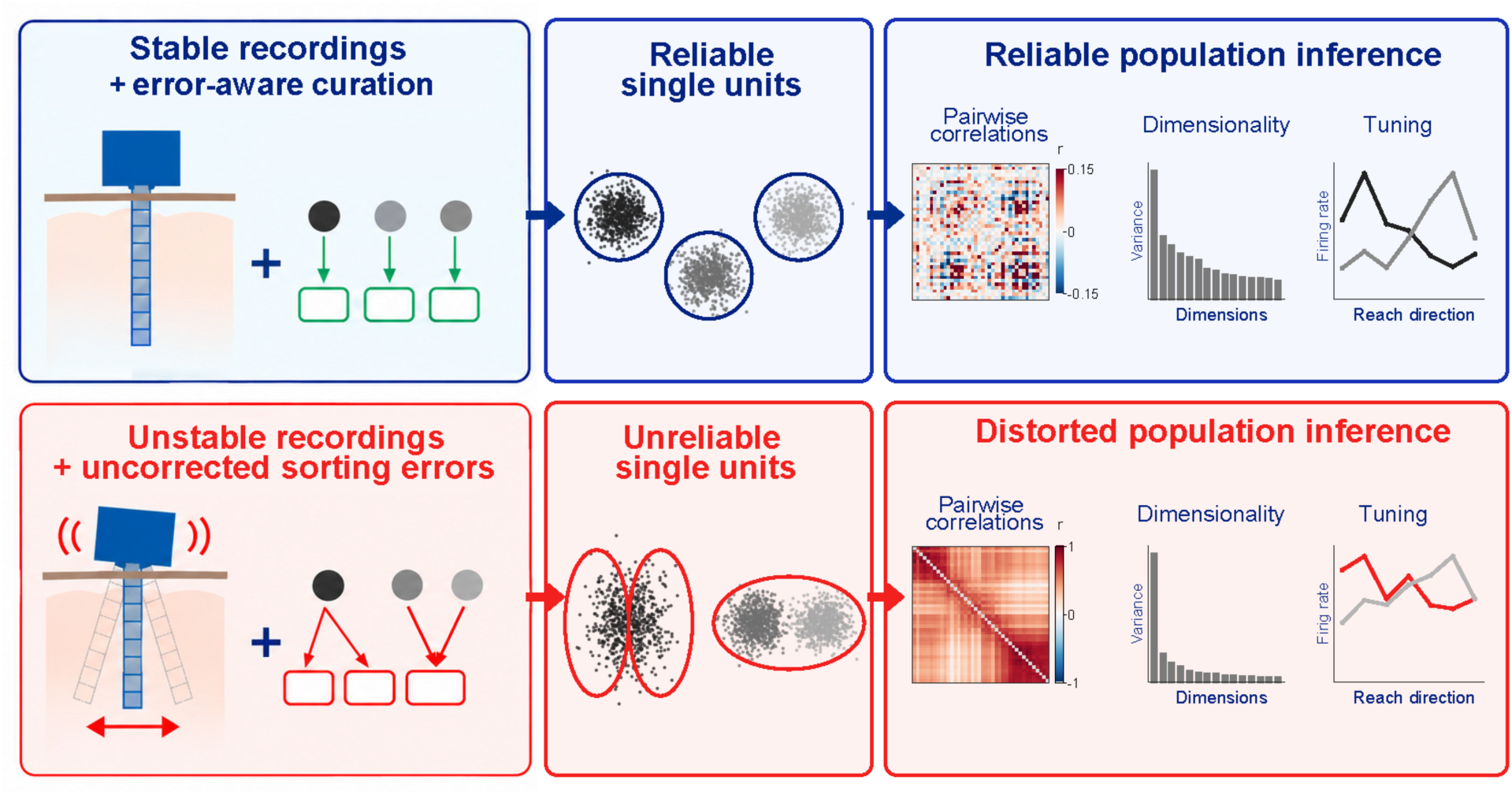

## Introduction

Recent advances in electrophysiological recording technologies have led to the development of high-channel-count, high-density probes such as Neuropixels^1,2^ and Neuroscroll^3^. These probes feature a laminar architecture with densely packed recording sites enabling neural population recordings from up to hundreds of neurons per shank. The compact footprint of high-density probes permits simultaneous deployment of multiple probes for investigating multi-regional neural population dynamics. As a result, large-scale, long-duration, multi-regional experiments are becoming routine in rodent studies^4–8^. Scaling recordings across more brain regions and longer durations will enable rigorous investigations of distributed neural mechanisms underlying complex behaviors.

Recent work in non-human primates (NHPs) has introduced the Neuropixels 1.0 NHP probe for high-density neural population recordings ^9^. The NHP probe is almost identical to the rodent electrode and was revised to aid penetration of the meninges. Thus, Neuropixels 1.0 NHP probe promises to deliver experimental scaling in primates, with their long lifespans and complex brain architecture, as routinely as in rodents. However, existing drive systems remain bulky, and cannot be implanted ^9,10^. Current approaches employ hydraulic microdrives or rail-based systems and so require head fixation for daily insertion. These approaches are well suited for conventional microelectrode recordings but become difficult to scale when deploying multiple silicon probes. A compact screw-driven actuation mechanism can provide independent probe advancement ^11,12^ without bulky hydraulic or motorized components, enabling multiple Neuropixels probes to be packaged within an implantable drive. This compactness is critical for scaling both probe number and recording duration, including chronic and ambulatory, head-free experiments.

Recording stability presents another challenge in primates, likely stemming from the size and weight of monkeys and humans compared to rats and mice. Without proper stabilization, probes are susceptible to fast-changing movement artifacts from cardiac pulsation, respiration, and other mechanical sources^13^. Similar issues are observed in acute human intraoperative studies^14,15^. Careful tissue preparation and mechanical stability of the apparatus housing the probe can substantially minimize these fast-movement artifacts ^9,14,16^, reducing dependence on specialized motion-correction algorithms^13,17^. Slow drift, arising from gradual tissue relaxation after probe insertion, is harder to avoid. Previous NHP Neuropixels studies have reported slow drift on the order of 15 µm^13,18–20^. Such displacement can alter the relationship between neurons and recording sites over longer recordings. Maintaining mechanical stability is therefore essential for preserving reliable single-unit isolation as recording duration increases.

Scaling multi-regional investigations also requires automated spike sorting and scalable post-sorting curation. Historically, spike sorting and curation have relied on manual or semi-manual approaches^21–25^. While effective for small datasets, manual and semi-manual approaches are labor-intensive, prone to human error, and fundamentally non-scalable ^26^. Large-scale, high-density datasets therefore require automated spike-sorting algorithms ^20,27–32^. Kilosort, a widely used automated spike-sorter, performs well in detecting units ^20,27,28,33^ but its cluster labels do not by themselves establish reliable single-unit identity.

Automated spike sorting often introduces characteristic errors in single-unit identification ^28,34^. Errors in single-unit identification can distort spike-spike relationships, alter estimates of population structure, and misrepresent the tuning and temporal dynamics of individual neurons ^35–38^. These biases can consequently distort inferences about neural population organization and interactions. Reliable single-unit identification at scale therefore requires post-sorting curation that can detect characteristic spike-sorting errors rather than relying on automated cluster labels alone. Importantly, spike-sorting errors are not corrected by recording more neurons or collecting more behavioral trials ^37,38^.

Here, we present an integrated hardware–software solution for scalable, high-density, multi-regional electrophysiology in NHPs. The hardware component, the Multi-regional Implantable NeuroDrive (MIND), is a compact implantable microdrive that supports stable recordings from independently positioned Neuropixels probes. The software component, UnitAssay, is an automated post-sorting curation framework that first evaluates cluster isolation and then tests for characteristic spike-sorting errors. We demonstrate the platform across 147 recording sessions and 53,346 Kilosort 4 clusters from two rhesus macaques performing reaching and eye-movement tasks with recordings spanning parietal, prefrontal, and motor cortices. Together, MIND and UnitAssay provide an integrated framework for scaling recordings across brain regions, recording duration, and neuronal populations while supporting reliable single-unit identification to investigate distributed neural mechanisms underlying complex behavior in NHPs.

## Results

### Multi-regional Implantable NeuroDrive (MIND)

We developed MIND, an implantable microdrive for multi-regional high-density neural population recordings in non-human primates using 10 mm Neuropixels 1.0 NHP probes (**Fig 1**). A key feature of MIND is the use of a screw-driven insertion mechanism, which replaces bulky hydraulic or motorized systems (**Fig 1A**). A precision leadscrew engages with a dovetail shuttle that securely anchors the Neuropixels base via a microscrew, enabling smooth and controlled advancement of the probe into brain tissue. Screw-driven probe insertion enables precise depth control and is central to scaling Neuropixels probes in count. Scaling stems from compact packaging of multiple probes within an 18 mm footprint (**Fig 1B**). The actuator block (23 mm height and 18 mm diameter) houses three probes in MIND-3 configuration and can be readily scaled to MIND-6, which accommodates up to six Neuropixels probes (**Fig 1C**). The compact geometry of the actuator block further enables simultaneous use of two drives when targeting distant brain regions.

**Figure 1.**
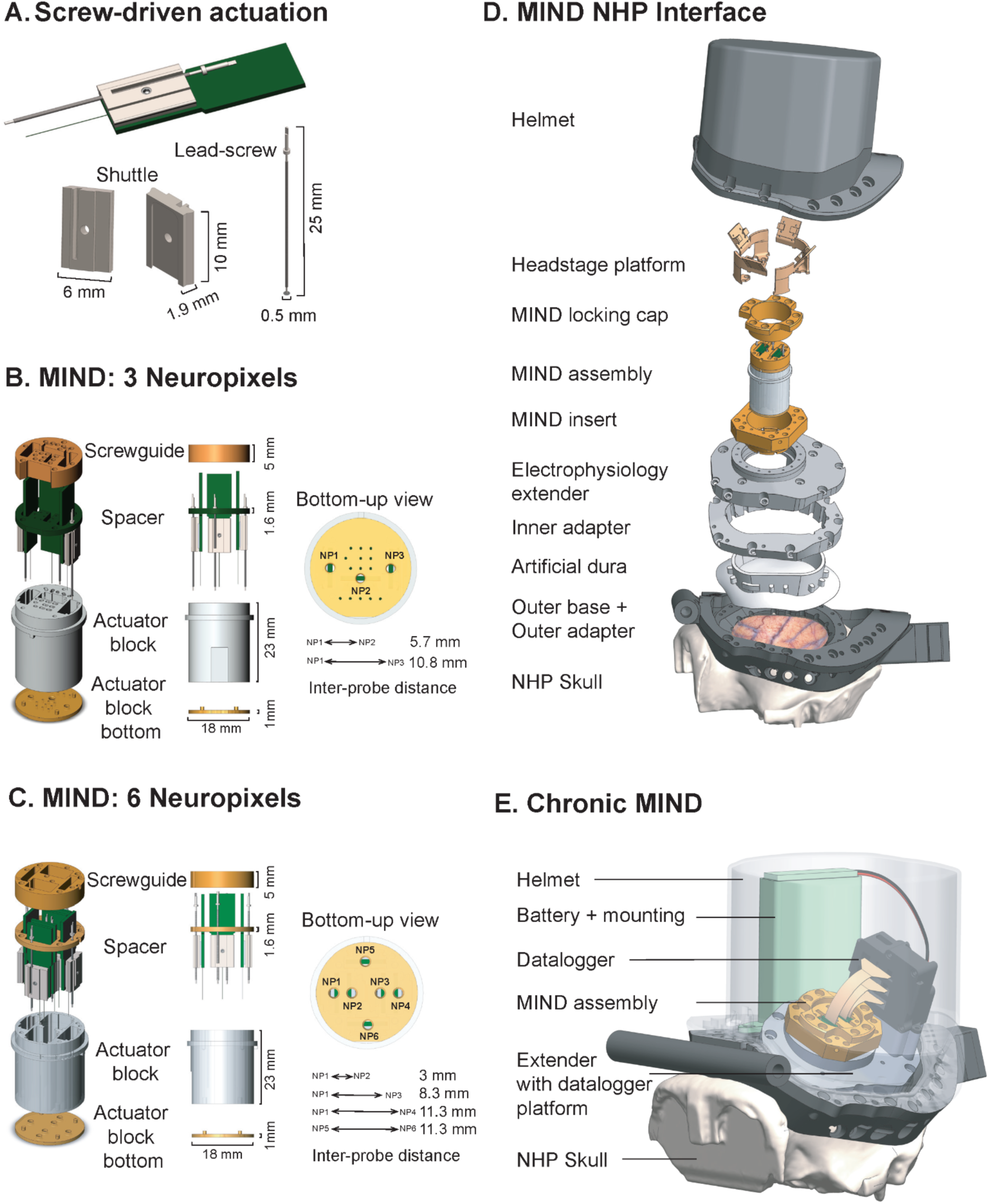
MIND: A scalable microdrive for multi-regional, high-density recordings in non-human primates. **(A)** Screw-driven actuation mechanism showing the leadscrew and dovetail shuttle assembly that enable precise, controlled probe advancement. **(B)** MIND-3 configuration housing three Neuropixels 1.0 NHP probes within an 18 mm diameter actuator block (23 mm height). Spherical disc at the actuator block base interfaces with meninges to minimize motion artifacts. Bottom-up view shows probe arrangement with inter-probe distances of 5.7-10.8 mm. Small holes in the actuator block accommodate microelectrodes and optrodes for electrical stimulation and optogenetics experiments. **(C)** MIND-6 configuration accommodates six probes within the same compact footprint of 18 mm. Bottom-up view shows probe spacing ranging from 3-11.3 mm for flexible multi-regional targeting. **(D)** Complete MIND-NHP interface system. The exploded view shows the modular implant architecture from top to bottom: protective helmet, headstage platform, MIND locking cap, MIND assembly, MIND insert, electrophysiology extender, inner adapter, artificial dura, outer base with outer adapter, and custom skull interface. **(E)** Chronic MIND configuration for chronic and ambulatory, head-free recordings. Integrated datalogger and battery system within protective helmet enable overnight and multi-day unrestrained recordings during in-cage behavior.

At its base, the actuator block is sealed by a spherical disc (1 mm thick, 18 mm diameter) that interfaces directly with the meninges. In our observation, the actuator block bottom proved critical for stable neural recordings by minimizing fast movement artifacts arising from cardiac pulsation, respiration, and other mechanical sources. The MIND interfaces with a custom implant insert that secures the drive to an electrophysiology extender, allowing flexible targeting of distinct brain-regions. The extender mates with the cranial chamber, providing a stable and accessible window to the craniotomy and underlying tissue (**Fig 1D**). The entire drive-assembly is enclosed within a protective *helmet,* crucial to scaling-up in duration. When paired with a datalogger (**Fig 1E**), MIND supports chronic recordings and head-free recordings, enabling unrestrained and in-cage experiments.

### Multi-regional multi-probe recordings

We used MIND-3 and MIND-6 to perform 38 Neuropixels insertions in adult rhesus macaque monkeys across 20 experimental days. To evaluate MIND’s performance across single and multi-region configurations, we deployed single (9 days), two (5 days), three (5 days), or four (1 day) 10 mm Neuropixels probes targeting parietal, prefrontal, premotor and motor cortices (**Fig 2A**). Each experimental day comprised a median of six (Min: 1, Max: 11) recording sessions per probe. This yielded a total of 147 recording sessions (2 -36.5 mins each) across all monkeys, probes and experimental days. Figure 2 shows an example recording from premotor cortex (PMd activity) during a center-out reaching task requiring reaches to one of seven randomly cued targets. Single-neuron (**Fig 2B**) and population-level (**Fig 2C**) recordings revealed diverse temporal dynamics around reach onset, with neurons exhibiting distinct activity profiles during planning and execution phases. Examining recordings across the depth of the probe revealed structured temporal patterns during reach initiation, demonstrating coordinated activity across local PMd populations. Depth dependent changes in firing activity illustrate the advantage of high-density sampling across cortical depths.

**Figure 2.**
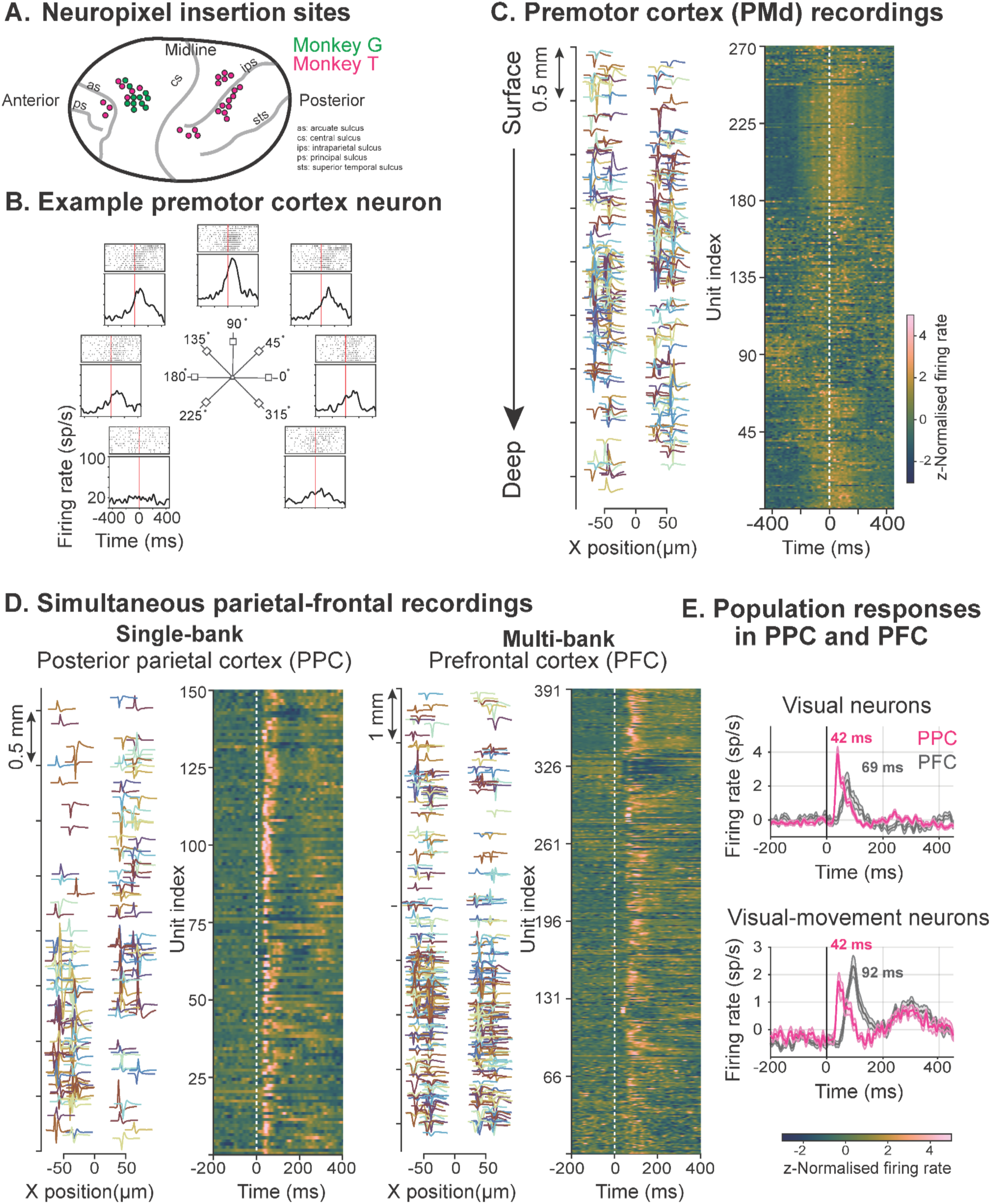
Multi-regional high-density recordings. **(A)** Recording locations across rhesus macaque cortex over 20 experimental days using 1-4 Neuropixels probes (single: 9 days, two: 5 days, three: 5 days, four: 1 day). **(B)** Example PMd neuron during center-out reaching to seven targets. Peri-event histograms and rasters aligned to reach onset (0 ms) show directionally tuned responses, with peak activity for the 90° target direction. **(C)** PMd population activity across cortical depth during center-out reaching for an example recording session. Left: Spike waveforms for 270 Kilosort 4-detected clusters spanning cortical depth (Bank 0, 3.84 mm). Right: Population firing rate heatmap for the same 270 clusters aligned to reach onset (white dashed line, 0 ms). Clusters sorted by depth. Color indicates z-normalized firing rate, revealing structured temporal patterns during reach initiation. **(D)** Simultaneous posterior parietal cortex (PPC) and prefrontal cortex (PFC) recordings during two-target free-choice eye-movement task, using two MIND drives. Left: PPC population activity (150 clusters) recorded from a single bank (Bank 0). Right: PFC population activity (391 clusters) across two banks (Banks 0 and 1; spanning 7.68 mm), with channels selected using a classification-based algorithm. Both regions show increased firing following visual target onset (0 ms, white dashed line). Units sorted by depth. **(E)** Population responses of curated single units reveal hierarchical processing between PPC and PFC. Top: Average firing rates of visually responsive neurons (PPC: n=19, magenta; PFC: n=12, gray) show increased activity following target onset. Bottom: Visual-movement neurons (PPC: n=20; PFC: n=14) respond during both target presentation and saccade execution. PFC neurons exhibit delayed responses relative to PPC neurons. Shaded regions indicate SEM.

To test the scaling capability of MIND, we simultaneously deployed two MIND drives targeting spatially separated parietal and prefrontal cortices, during a two-target free-choice eye-movement task (**Fig 2D**). The compact geometry of MIND enabled this dual-drive configuration. Each drive contained a single Neuropixels probe targeting lateral intraparietal area (LIP) in posterior parietal cortex and frontal eye field (FEF) in prefrontal cortex. In LIP, neural activity was concentrated in the deepest bank (Bank 0), and all channels within that bank were recorded. In FEF, neural activity extended across multiple banks. To increase single-unit yield across a larger probe depth, we selected channels across two banks (Banks 0 and 1, spanning 7.68 mm), using a classification-based selection algorithm that identifies high-quality channels across non-contiguous probe regions^39^. Population activity in both regions showed increased firing following target onset (**Fig 2E**). FEF neurons showed delayed responses relative to LIP neurons, consistent with hierarchical visuomotor processing. Together, these experiments demonstrate that MIND can scale recordings both across distant brain regions through simultaneous deployment of multiple drives and across cortical depth through multi-bank channel selection.

### MIND enables stable recordings

We next evaluated the stability of MIND by examining the displacement of each Neuropixels probe relative to brain tissue. To quantify probe displacement, we used Kilosort 4 (KS4) drift estimates and calculated the 5th-to-95th percentile displacement range (range₉₀) for each recording session (**Fig. 3A, B**; Methods). Across 137 artifact-free recording sessions, median drift was 5.2 µm, and 131 of 137 sessions (96%) remained below 20 µm, corresponding to one row spacing on the Neuropixels 1.0 NHP probe (**Fig. 3B–E**). Recordings remained stable across multiple sessions collected throughout an experimental day, including intervening breaks during which the MIND remained implanted (**Fig. 3A, B,F,G**). A subset of recording sessions with displacement artifacts showed much larger and sometimes coordinated movements across probes; an example is shown in Fig. 3H.

**Figure 3.**
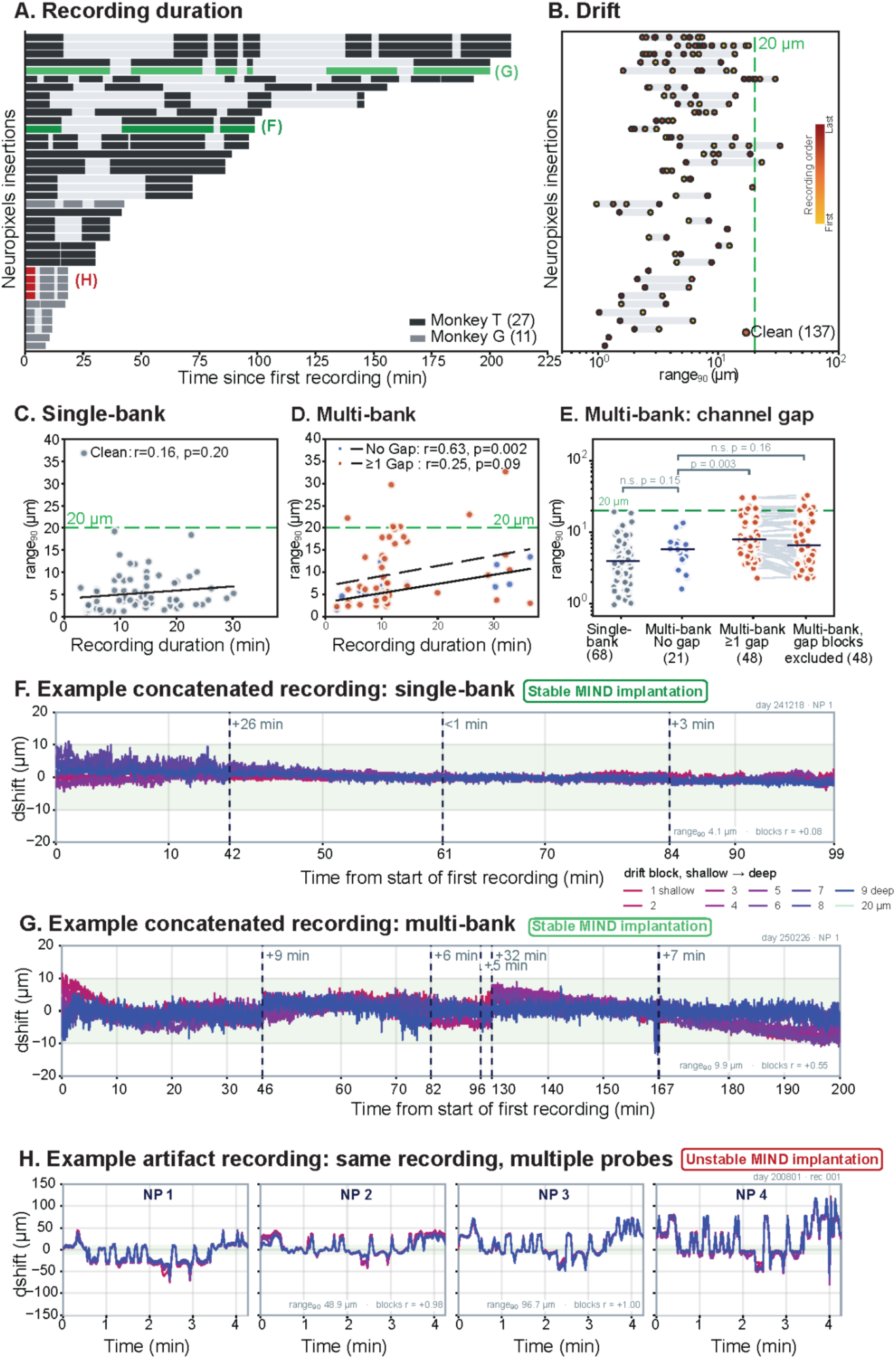
MIND enables stable Neuropixels recordings. **(A)** Recording timelines across 38 Neuropixels probe insertions in Monkey T (n = 27) and Monkey G (n = 11). Horizontal bars indicate individual recording sessions positioned relative to the start of the first recording for each insertion. Highlighted recordings correspond to the examples shown in panels F–H. **(B)** Probe displacement across 137 artifact-free recording sessions corresponding to the Neuropixels probe insertions shown in A. Each point shows the 5th-to-95th percentile displacement range (range₉₀) for one session, colored by recording order. The green dashed line indicates the 20 µm spacing between Neuropixels recording-site rows. **(C)** Probe displacement as a function of recording duration for 68 artifact-free single-bank sessions. The black line shows the linear fit. **(D)** Probe displacement as a function of recording duration for artifact-free multi-bank sessions. In non-contiguous multi-bank configurations, some Kilosort depth blocks can span an unrecorded gap between recorded channel groups. Blue points indicate recordings without gap-spanning depth blocks (n = 21); orange points indicate recordings containing at least one gap-spanning block (n = 48). Solid and dashed lines show the corresponding linear fits. **(E)** Displacement estimates for single-bank and multi-bank recordings with and without gap-spanning depth blocks. For recordings with gap-spanning blocks, displacement is also shown after excluding those blocks. Points indicate recording sessions, horizontal bars indicate medians, and gray lines connect paired estimates. **(F)** Example concatenated single-bank recording from a stable MIND implantation. Colored traces show Kilosort displacement estimates across nine depth blocks, ordered from shallow to deep. Vertical dashed lines indicate transitions between recording blocks; labels indicate the interval between recordings. **(G)** Example concatenated multi-bank recording from a stable MIND implantation, shown as in F. **(H)** Example recording with displacement artifacts obtained simultaneously from four Neuropixels probes positioned using MIND. Displacement traces are shown for each probe. For panels F–H, range₉₀ and correlations between depth-block displacement traces are shown within each panel.

Because KS4 drift estimation is affected by non-contiguous channel configurations, we separately examined single-bank and multi-bank recordings. Single-bank recordings provide the most direct estimate of probe stability because all recorded channels are spatially contiguous. Across 68 artifact-free single-bank recordings, median drift was 4.0 µm (range, 1–19.2 µm; recording duration, 2.9–30.2 min; **Fig. 3C**). Probe displacement did not significantly increase with recording duration (r = 0.16, p = 0.20, Pearson correlation), indicating little accumulation of displacement over the duration of these recordings. In multi-bank recordings, KS4 depth blocks can span gaps between recorded banks and combine signals from spatially separated contacts, inflating estimates of displacement. Multi-bank recordings without channel gaps were stable and demonstrated a median drift of 5.8 µm (range, 1.6–13.4 µm), whereas recordings with one or more gap-spanning blocks had a median drift of 7.9 µm (range, 2.3–30.3 µm; **Fig. 3D, E**). Excluding gap-spanning blocks reduced the estimated displacement (**Fig. 3E**), indicating that part of the apparent increase in multi-bank recordings reflects the limitations of KS4 drift estimation rather than probe instability. Together, these results demonstrate that MIND provides stable Neuropixels probe implantation across recording configurations and recording duration.

### Spike-sorting errors limit reliable single-unit identification

We next examined the performance of Kilosort 4 (KS4), an automated spike sorter in its ability to reliably identify single units. Examination of KS4 output revealed several characteristic spike-sorting errors, including over-splitting, over-merging, duplicate detections, and intermittent unit presence (**Fig. 4**). In over-splitting, spikes from a single neuron are erroneously assigned to multiple clusters. These clusters typically show overlap or close proximity in feature space. Their spike trains will show temporal relationships consistent with a single underlying neuron, as revealed by cross-correlogram analysis (**Fig. 4A**). Importantly, waveforms may or may not be similar, as a waveform shape can vary with the distance between a neuron and the recording site. Over-merging occurs when spikes from distinct neurons are grouped into a single cluster. Feature-space analysis can reveal separate spike populations within the cluster, suggesting that spikes from multiple neurons have been combined. Temporal measures such as refractory-period violations may provide additional evidence for over-merging but are not necessarily present (**Fig. 4B**). Duplicate detection, or re-detection, occurs when the same spike events are represented in more than one cluster. During template matching, Kilosort fits templates to the raw voltage and subtracts the fitted activity from the signal. Residual activity from an incompletely subtracted spike can subsequently be fit by another template, which can result in re-detection of the same spike event. Duplicate clusters therefore produce highly coincident spike trains, often with a consistent small temporal offset that can be identified by cross-correlogram analysis (**Fig. 4C**). Intermittent presence occurs when spikes are assigned to a cluster during only part of the recording, resulting in unstable single-unit isolation over time (**Fig. 4D**). These errors have distinct signatures across waveform, feature-space, correlogram, and temporal analyses, consequently these cannot be reliably identified using a single conventional quality metric. Manual curation can identify some of these errors but is subjective and difficult to scale to the hundreds of clusters generated in each high-density recording session.

**Figure 4.**
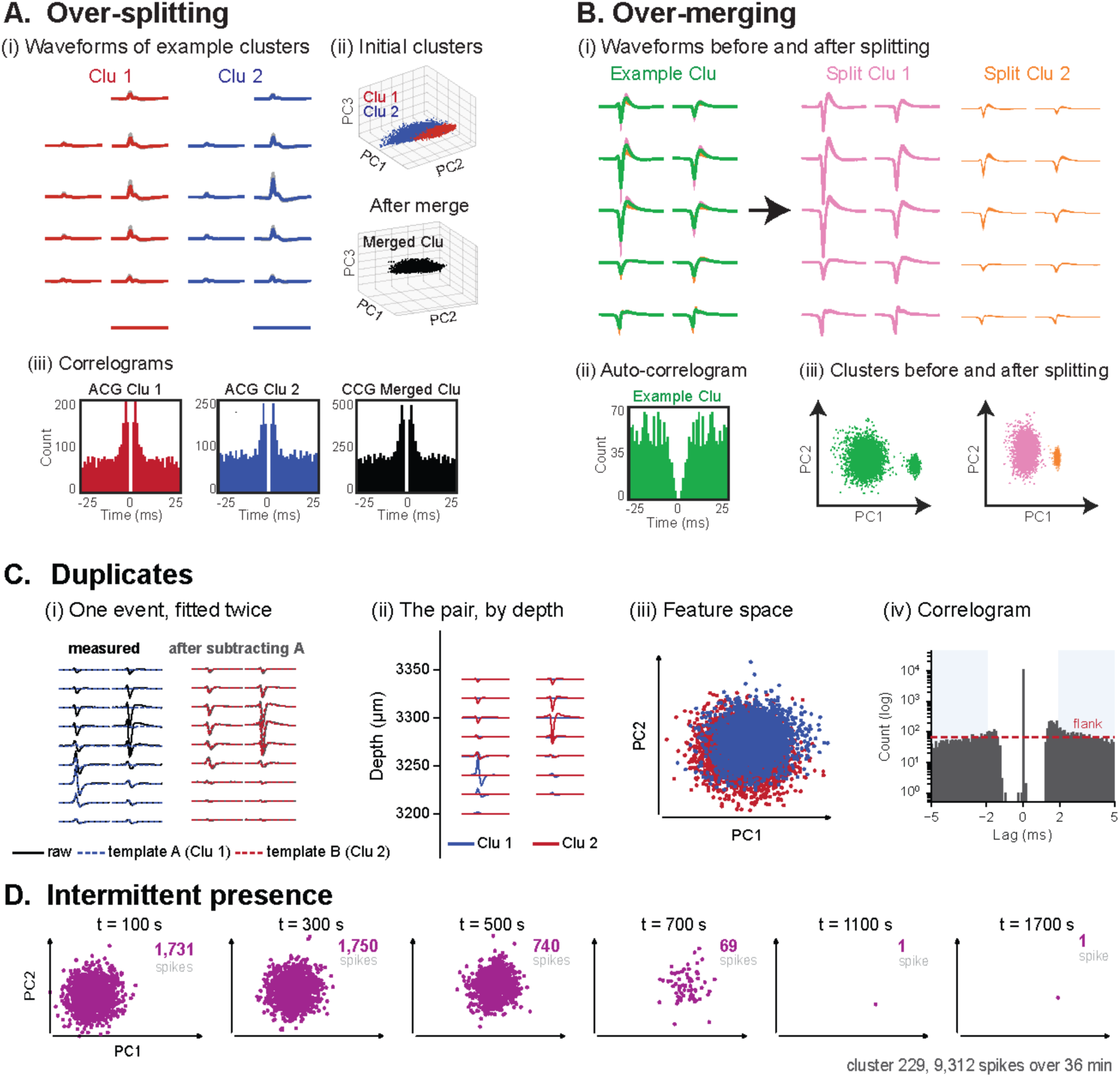
Four characteristic errors in Kilosort 4 automated spike-sorting output. (A) Over-splitting: spikes from a single neuron are partitioned across multiple clusters. (i) Example waveforms from two clusters (red and blue) across channels. (ii) Principal-component feature space of the two clusters before merging (top) and after merging into a single cluster (bottom). (iii) Autocorrelograms (ACGs) of the individual clusters and the cross-correlogram (CCG) between the two clusters; CCG provides evidence that the two clusters originate from the same neuron. **(B) Over-merging:** spikes from distinct neurons are grouped into a single cluster. (i) Waveforms of the original cluster (green), showing separation into two distinct waveform populations after splitting (pink and orange). (ii) ACG of the original cluster can retain a clean refractory period despite containing spikes from multiple neurons. (iii) Principal-component feature space shows two separable populations within the original cluster, which are assigned to distinct clusters after splitting **(C) Duplicate detection (re-detection):** the same spike event is detected more than once and assigned to separate clusters. (i) Example raw voltage for a shared event and the residual after subtracting the fitted template of cluster 1, revealing the waveform assigned to cluster 2. (ii) Waveforms of the two clusters across recording depth. (iii) The two clusters overlap in feature space. (iv) Cross-correlogram between the two clusters; the peak occurs at a non-zero lag. **(D) Intermittent presence:** a cluster is present for only part of the recording. Principal-component feature space is shown at six time points across the session, with the number of spikes indicated at each time point. The example cluster contained 9,312 spikes over a 36-min recording.

To reliably identify single units, scalable software is needed that can curate the recordings obtained with MIND. Developing such software requires an understanding of the spike-sorting errors present in the MIND recordings and an independent ground truth against which they can be evaluated. We therefore generated synthetic Neuropixels recordings using the MEArec Python package through SpikeInterface ^28,40^, with known ground-truth neuron identities and spike times. Importantly, the simulations were designed to reflect properties of our experimental recordings, using Neuropixels geometry and varying neuronal density, recording duration, noise, drift, bursting, and firing-rate statistics (see **Supp. Fig. 1A–D, Methods**). We found that synthetic data was important for defining and validating the distinction between duplicates and over-splitting. An over-split fragment can represent either a divided portion of a neuron’s spikes or a re-detected copy of spikes that are also retained in another cluster (**Supp. Fig. 1E**). Both duplicate and over-splitting errors can alter spike-spike relationships and lead to erroneous estimates of relationships between neurons (**Fig. 5D**). In addition, duplicate units introduced redundancy and consequently reduced estimated dimensionality (**Fig. 5E**). These synthetic ground-truth results validate the impact of spike-sorting errors on subsequent neural data analysis and highlight the importance of reliable post-sorting curation software.

**Figure 5.**
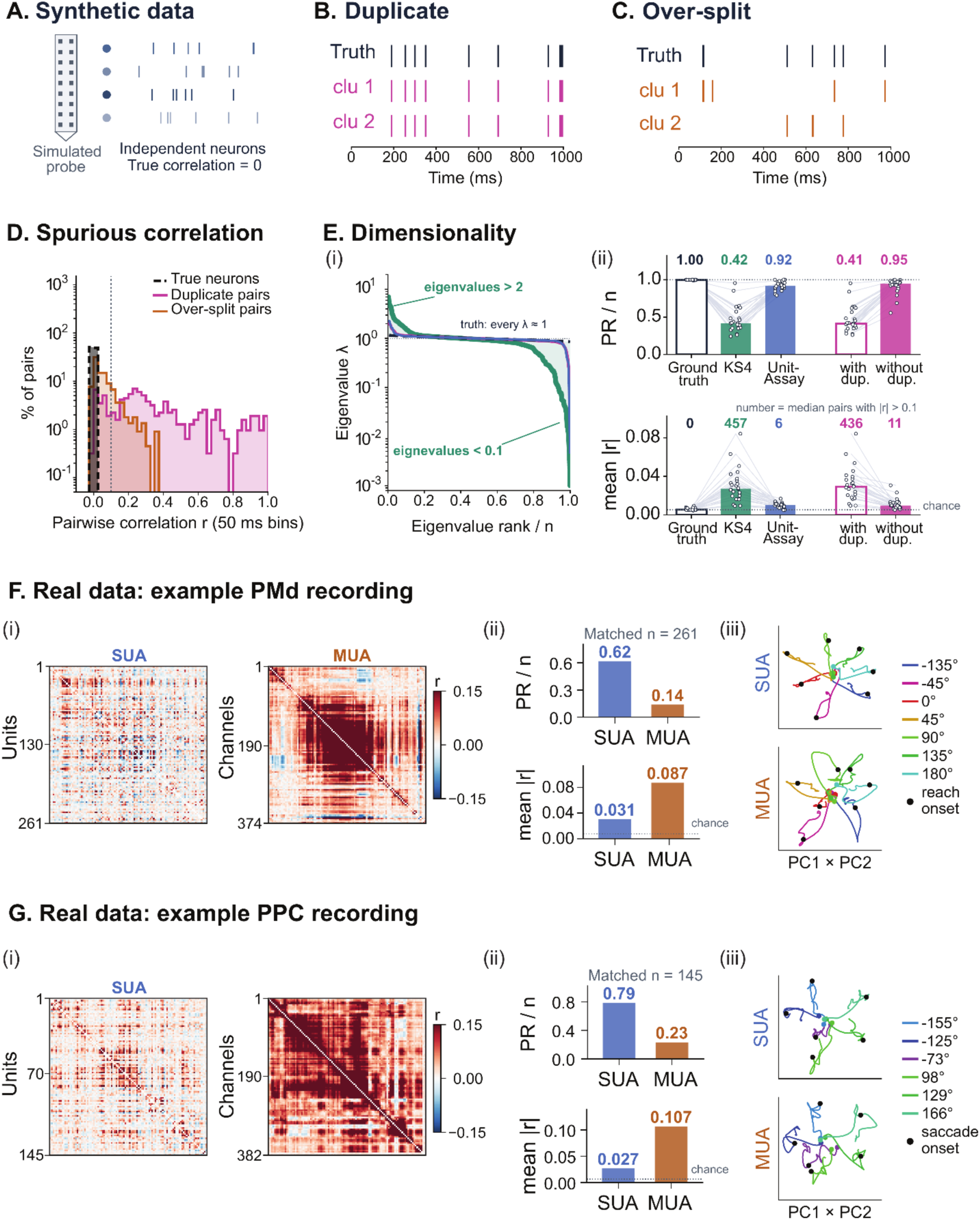
Spike-sorting errors distort neural correlations and population dimensionality. **(A)** Synthetic Neuropixels recordings generated from independent neurons with zero ground-truth pairwise correlation. **(B)** Example KS4 duplicate error. A single ground-truth spike train is detected twice and assigned to two clusters, producing redundant spike trains. **(C)** Example KS4 over-split error. Spikes from a single ground-truth neuron are divided between two clusters. **(D)** Distribution of pairwise spike-count correlations calculated in 50-ms bins for ground-truth neuron pairs, duplicate pairs, and over-split pairs. Duplicate and over-split errors introduce correlations that are absent between the underlying independent neurons, with duplicates producing the strongest spurious correlations. **(E)** Effect of sorting errors on population dimensionality. **(i)** Normalized eigenvalue spectra for ground truth and spike-sorted populations. **(ii)** Normalized participation ratio (PR/n; top) and mean absolute pairwise correlation, |r| (bottom), for ground truth, Kilosort 4 output, UnitAssay-curated output, and Kilosort 4 output evaluated before and after removal of duplicate units. Points indicate individual simulated recordings and lines connect matched recordings. **(F)** Comparison of UnitAssay-curated single-unit activity (SUA) and multi-unit activity (MUA) in an example premotor cortex (PMd) recording during a center-out reaching task. **(i)** Pairwise spike-count correlation matrices for SUA and MUA populations. **(ii)** Normalized participation ratio (PR/n; top), calculated at matched population sizes, and mean absolute pairwise correlation (bottom), calculated across all available units or channels. **(iii)** Reach-direction-specific neural population trajectories projected onto the first two principal components (PC1 × PC2) for SUA and MUA. Colors indicate reach directions, and black dots mark reach onset. **(G)** Same analysis as in (F) for an example posterior parietal cortex (PPC) recording during a luminance-selection task, comparing UnitAssay-curated SUA with MUA. **(i)** Pairwise spike-count correlation matrices for SUA and MUA populations. **(ii)** Normalized participation ratio (top) and mean absolute pairwise correlation (bottom). **(iii)** Saccade-direction-specific neural population trajectories projected onto the first two principal components for SUA and MUA. Colors indicate saccade directions, and black dots mark saccade onset.

We next examined how single unit curation impacts estimates of population correlations, dimensionality, and neural dynamics in experimental recordings. We compared the single-unit activity (SUA), obtained after curating spike-sorting outputs, with multi-unit activity (MUA), obtained from threshold crossings on individual Neuropixels channels. We analysed recordings from two different areas: premotor cortex (PMd) during a center-out reaching task and posterior parietal cortex (PPC) during a luminance-selection task (**Fig. 5F, G**). In both brain regions, mean absolute pairwise correlations were higher for MUA than SUA (PMd: 0.087 vs. 0.031; PPC: 0.107 vs. 0.027; **Fig. 5Fii, Gii**). MUA exhibited reduced population dimensionality (PR/n) compared with SUA (PMd: 0.14 vs. 0.62; PPC: 0.23 vs. 0.79; **Fig. 5Fii, Gii**). The low-dimensional neural trajectories derived from MUA differed from those derived from SUA and captured less direction-specific population structure (cross-validated R² values: PMd, MUA = 0.52 vs. SUA reliability ceiling = 0.81; PPC, MUA = 0.40 vs. SUA reliability ceiling = 0.58; see **Methods**). These results demonstrate that threshold-crossing MUA does not faithfully recover all aspects of neural population dynamics captured by curated SUA.

### UnitAssay enables automated post-sorting curation

UnitAssay is automated software for curating spike-sorting output (**Fig. 6A**). UnitAssay consists of two stages. Layer 1 (screen) evaluates per-cluster isolation using L-ratio and its stability over time. Layer 2 (curate) applies a decision flow comprising five tests and assigns a curation outcome to each cluster. Clusters that fail curation are subsequently assigned to one of four characteristic error classes: over-split, over-merged, duplicate, or intermittent.

**Figure 6.**
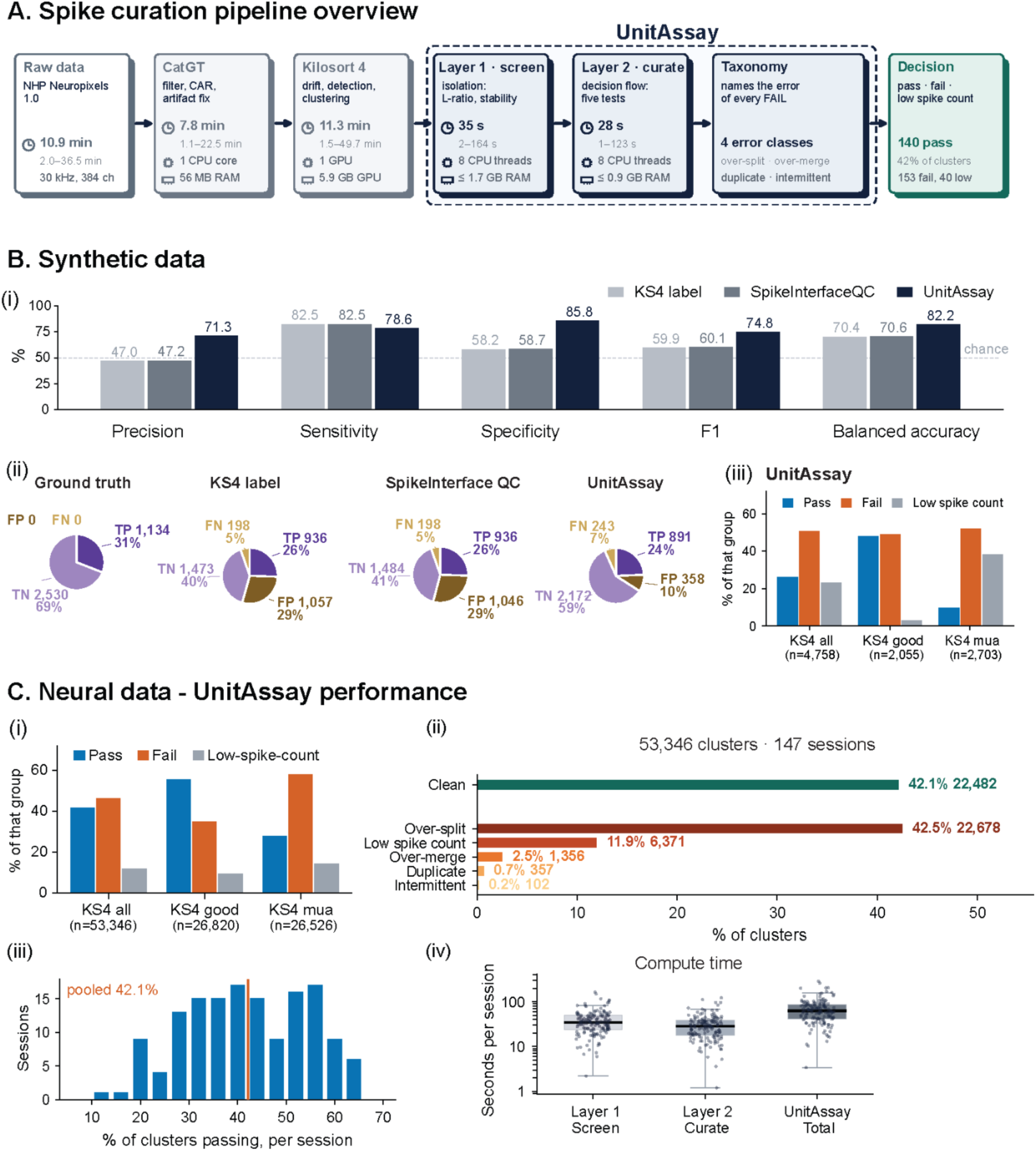
UnitAssay pipeline and performance on synthetic ground-truth and neural datasets. **(A)** Overview of the spike-sorting and curation pipeline. Raw Neuropixels recordings are preprocessed with CatGT and sorted with Kilosort 4. UnitAssay then operates in two stages: Layer 1 screens cluster isolation and isolation stability, and Layer 2 applies the sequential curation decision flow. A subsequent taxonomy step assigns an error class to failed clusters. The final output assigns each cluster a pass, fail, or low-spike-count outcome. Processing time and computational requirements are shown for each stage. **(B)** Performance on synthetic recordings with known ground-truth neuron identities. **(i)** Precision, sensitivity, specificity, F1 score, and balanced accuracy for Kilosort 4 labels, standard SpikeInterface quality control, and UnitAssay. **(ii)** Confusion-matrix composition for ground truth, Kilosort 4 labels, SpikeInterface quality control, and UnitAssay. TP, true positive; TN, true negative; FP, false positive; FN, false negative. **(iii)** UnitAssay pass, fail, and low-spike-count outcomes for all synthetic Kilosort 4 clusters and separately for clusters labeled “Good” or “MUA”. **(C)** UnitAssay outcomes on neural recordings. **(i)** Distribution of pass, fail, and low-spike-count outcomes across all 53,346 Kilosort 4 clusters and separately for clusters labeled “Good” or “MUA.” **(ii)** Final UnitAssay classification across 147 recording sessions. **(iii)** Distribution of the percentage of clusters passing curation across recording sessions; the vertical line indicates the pooled pass rate. **(iv)** UnitAssay computation time for Layer 1, Layer 2 per recording session.

To evaluate UnitAssay performance, we tested it against synthetic ground-truth data (see **Methods**). UnitAssay outperformed both Kilosort 4 labels and standard SpikeInterface quality control, with the largest improvement in specificity (**Fig. 6B**). Specificity increased to 85.8%, compared with 58.2% for Kilosort 4 and 58.7% for SpikeInterface quality control. The increase in specificity was due to improved rejection of clusters that did not correspond to clean single units, with more true negatives and fewer false positives (**Fig. 6B ii**). UnitAssay also showed higher precision, while sensitivity remained similar to KS4 and SpikeInterface quality control (**Fig. 6B**).

We next applied UnitAssay to 53,346 clusters across 147 neural recording sessions (**Fig. 6C**). UnitAssay classified 42.1% of the clusters as clean, single units. Importantly, UnitAssay classification was not dependent on the original KS4 label (**Fig. 6Ci**). UnitAssay rejected 34.4% of KS4 ‘good’ clusters and passed 28% of KS4 ‘MUA’ clusters. The fraction of clusters passing UnitAssay varied across sessions (**Fig 6C iii**) and was lower across sessions with drift-related artifacts (28.7%; see **Methods**). UnitAssay classified 42.5% of clusters as over-split, 11.9% as low-spike-count, and 2.5% as over-merged, while duplicate and intermittent classifications were less frequent. Because over-splitting and duplicate re-detection can arise from the same underlying neuronal spike train, the distinction between them is more difficult to validate in neural recordings without ground truth. Layer 1 and Layer 2 required, a median of 34.7 s (range, 2.2– 163.9 s) and 28.3 s (range, 1.2–122.7 s) per recording session, respectively, on CPU alone (**Fig. 6Civ**; see **Methods**), making the framework fast and computationally lightweight. Together, these results show that UnitAssay improves single-unit identification against known ground truth and enables curation of high-density neural recordings at scale.

### UnitAssay screening and error-specific curation

To evaluate whether a cluster represents a reliably isolated single unit, Layer 1 (Screen) examines the cluster in feature space, where each spike is represented by waveform-derived principal-component features. For each cluster, UnitAssay assigns a unit channel and a spatially local reference channel. Because signals from multiple nearby neurons can be detected across overlapping recording sites, cluster isolation must be evaluated relative to local neuronal activity. The *unit-channel* is selected based on maximal waveform energy and spike representation and captures the strongest activity associated with the cluster, whereas the *reference-channel* provides a local comparison containing predominantly non-cluster activity (**Supp. Fig. 2A**; see **Methods**). We quantified the separation between activity observed at the unit and reference channels using L-ratio ^41,42^. L-ratio uses Mahalanobis distance in principal-component feature space to quantify overlap between the cluster and local non-cluster activity (**Supp. Fig. 2B**). Layer 1 further examines whether isolation is maintained throughout the recording by recomputing L-ratio in consecutive 100-s windows. At the 60% isolation-stability criterion, 57.5% of Kilosort “Good” clusters met the criterion compared with 37.9% of “MUA” clusters (**Supp. Fig. 2C, D**). Together, L-ratio and isolation stability allow Layer 1 to screen for clusters that are both well isolated and stable over time before they proceed to Layer 2 curation.

Curation is challenging as no single metric can reliably identify all characteristic spike-sorting errors. Different errors produce distinct signatures in temporal and feature space, and the same metric can be informative for one error but inconclusive for the other (**Fig. 7**). Layer 2 (curate) therefore evaluates each KS4 cluster through a sequential decision flow that integrates temporal and feature-space evidence (**Fig. 7A**). Each cluster passes through a fixed sequence of tests for identity, refractory-period violations, within-cluster composition, contamination, and separation from neighboring clusters.

**Figure 7.**
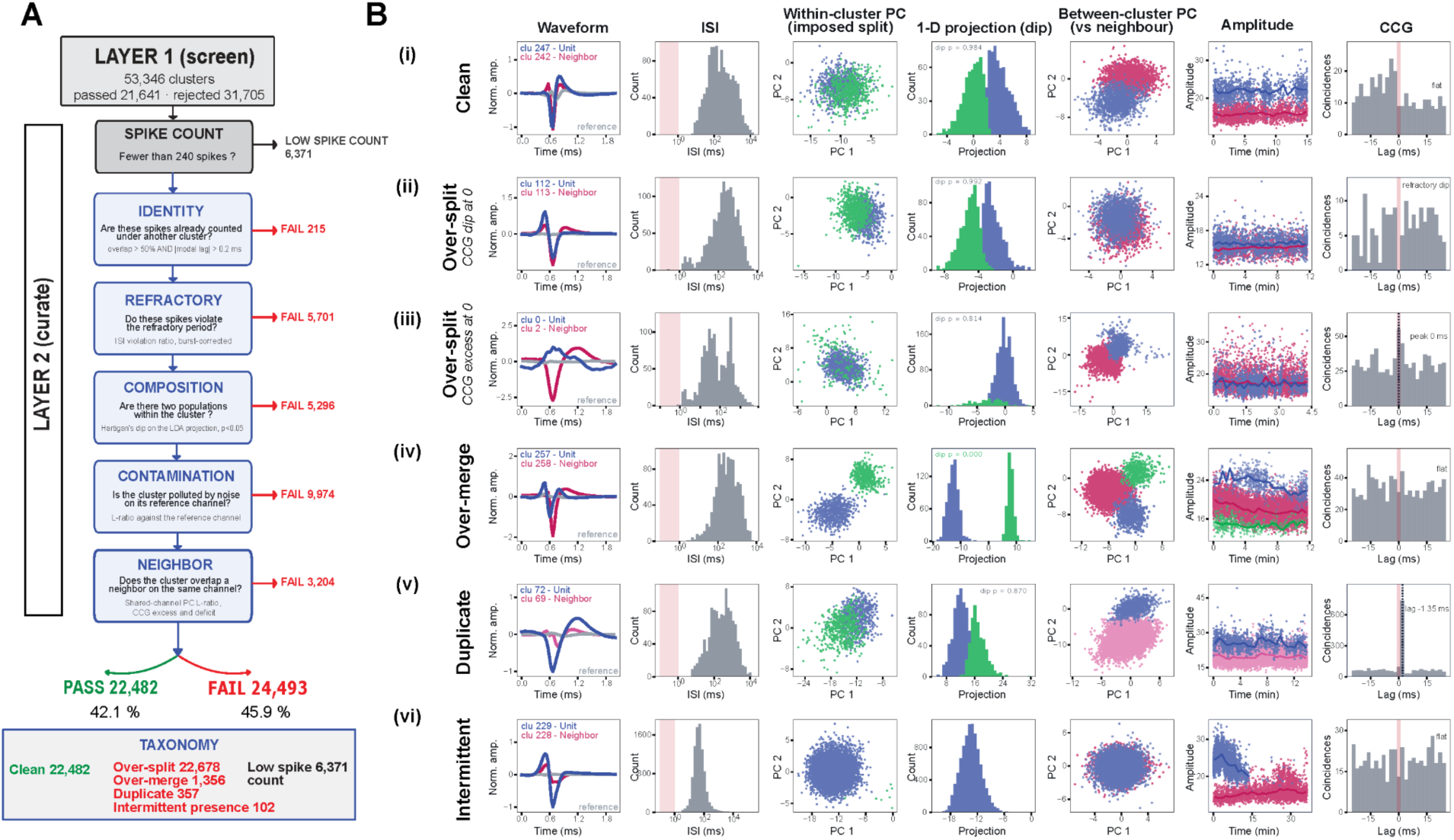
UnitAssay Layer 2 (Curate): sequential decision flow and error taxonomy. **(A)** Layer 2 evaluates each Kilosort 4 cluster using the Layer 1 screening outcome together with a fixed sequence of five tests. Clusters with fewer than 240 spikes are classified as low spike count and excluded from subsequent curation tests as the number of spikes is insufficient for reliable statistical evaluation. The identity test evaluates whether the same spikes have been detected under another cluster; the refractory test evaluates refractory-period violations after burst correction; the composition test evaluates whether more than one spike population is present within a cluster; the contamination test evaluates overlap with the local reference channel; and the neighbor test evaluates separation from nearby clusters using feature-space and temporal evidence. Clusters exit the decision flow when sufficient evidence determines a PASS or FAIL outcome. A subsequent taxonomy step classifies clusters that pass as clean and clusters that fail according to characteristic sorting errors. **(B)** Representative examples illustrating the measurements used during curation. Columns show waveform, inter-spike interval (ISI) distribution, within-cluster principal-component structure following an imposed split, one-dimensional projection used to assess bimodality, between-cluster principal-component space relative to a neighboring cluster, spike amplitude over time, and cross-correlogram (CCG). **(i) Clean:** the imposed split shows no significant bimodality, the unit remains separated from its neighboring cluster, and the CCG shows no prominent structure around zero lag. **(ii) Over-split, CCG gap at zero:** the unit and neighboring cluster overlap in feature space and their CCG shows a refractory-period gap around zero lag. **(iii) Over-split, CCG excess at zero:** the unit and neighboring cluster show overlapping feature-space structure, but the CCG instead shows increased coincident firing around zero lag. This example illustrates that CCG structure alone is insufficient to identify over-splitting and must be interpreted together with feature-space and other curation evidence. **(iv) Over-merged:** the imposed split reveals separable populations within the cluster, with significant bimodality in the one-dimensional projection. **(v) Duplicate:** the two clusters show overlapping feature-space structure and a sharp CCG peak at a consistent non-zero lag. **(vi) Intermittent presence:** spike amplitude and cluster representation change over the course of the recording.

The identity test evaluates re-detected duplicates by examining whether same spikes have been detected in another cluster based on shared spikes and their temporal offset. The identity test distinguishes between duplicates, zero-offset sharing, and other coincident activity (**Supp. Fig. 3**). The composition test evaluates whether a cluster contains multiple separable spike populations, as expected for over-merging, using complementary tests of bimodality and feature-space separation (**Supp. Fig. 4**). Over-splitting is identified from the pattern of evidence across the decision flow, including refractory-period violations, contamination relative to the local reference established in Layer 1, and separation from neighboring clusters in temporal and feature space. Examples in **Fig. 7B** illustrate that clean, over-split, over-merged, duplicate, and intermittent clusters are not distinguished consistently by any single metric. Notably, over-splitting itself can appear in different forms, with the two examples in **Fig. 7B** showing different cross-correlogram and feature-space signatures. Waveform shape, ISI distribution, within-cluster feature structure, neighboring-cluster separation, amplitude over time, and cross-correlogram structure contribute differently across error classes. This heterogeneity motivates the sequential decision flow rather than a single cluster-quality score. Final error classification is assigned in a separate taxonomy step after curation. Together, Layer 1 and Layer 2 screen and curate each KS4 cluster to identify reliably isolated single units.

## Discussion

Understanding neural-circuit mechanisms of complex behavior requires examining the activity of hundreds of individual neurons across brain regions in NHPs. Scaling electrophysiology requires solving two linked problems: implanting multiple high-density probes across brain regions and reliably identifying populations of individual neurons from the resulting recordings. Here, we address both challenges with an integrated hardware-software system. Multi-regional Implantable NeuroDrive (MIND) enabled stable, low-drift recordings from independently inserted Neuropixels probes across parietal, prefrontal, premotor, and motor cortices. UnitAssay enabled automated, reliable identification of single units while remaining fast and computationally lightweight. Using synthetic ground truth, we further show that errors in single-unit identification can distort estimates of neuronal interactions and population structure.

Scaling electrophysiology requires integrated hardware and software. The hardware must enable stable recordings from hundreds of individual neurons, while the software must ensure reliable identification of single units. These requirements are interdependent. Instability in recording hardware can compromise spike sorting and single-unit identification. Curation outcomes, in turn, provide a quantitative measure of recording quality and can inform the implantation protocol. MIND scales the recording hardware by housing up to six independently inserted Neuropixels probes within a compact drive. Recording scale can be increased further by deploying multiple drives simultaneously. MIND components are custom-designed and modular, enabling further scaling across probes and brain regions. UnitAssay reliably identifies single units, achieving 71.3% precision and 85.8% specificity against synthetic ground truth. UnitAssay is computationally efficient, requiring a median of 166 ms per cluster for curation. This makes UnitAssay well suited for increasingly large datasets as recording scale grows.

A key design feature of MIND is the integration of screw-driven actuation directly with each Neuropixels probe. A leadscrew coupled to the Neuropixels dovetail enables independent probe insertion and controlled advancement within a compact implantable geometry. Screw-driven microdrives have long been used in NHP electrophysiology for adjustable electrode insertion and high-resolution control of electrode movement ^43–45^. MIND adapts screw-driven actuation to high-density silicon probes within a compact implantable geometry. Multiple Neuropixels probes can therefore be positioned within a single 18-mm footprint and advanced independently. Other approaches have achieved dense multi-probe recordings using chamber-mounted linear rails coupled to external actuators ^9^. MIND instead incorporates independent probe actuation within the implant itself, avoiding the need for bulky external actuators for each probe. This compact geometry enables scaling while maintaining recording stability. Out of 137 artifact-free recording sessions, 131 showed probe drift below 20 µm, corresponding to one inter-row spacing between Neuropixels recording sites. This low drift is consistent with previous human and NHP Neuropixels studies highlighting probe-tissue motion and mechanical stabilization as important factors in recording stability ^9,15,46^.

Reliable single-unit identification requires validation against known ground truth ^33,40,47^. However, experimental ground truth does not scale to the hundreds of neurons targeted by high-density electrophysiology ^40,47^. Synthetic recordings provide a scalable alternative as the identity and spike times of every simulated neuron are known ^20,28,33,40^. Importantly, synthetic ground truth is critical for understanding the errors present in spike-sorted data and where they arise ^40^. Synthetic data can distinguish clusters affected by recording noise and contamination from errors introduced by the spike sorter, including over-merging, over-splitting and duplicate re-detection. Ground truth can further show why characteristic errors survive standard quality criteria and can be incorrectly identified as single units. This is important because different errors can produce similar signatures in downstream analyses. For example, shared artifacts, over-split fragments, and duplicate re-detections can all introduce structure in cross-correlograms that may be interpreted as neuronal interactions, including putative monosynaptic coupling (**Supp. Fig. 3**). Synthetic ground truth also enabled us to evaluate whether and how UnitAssay improved single-unit identification. UnitAssay increased specificity to 85.8%, compared with 58.2% for Kilosort 4 labels and 58.7% for standard SpikeInterface quality criteria, with a modest reduction in sensitivity. This gain in specificity reflects improved rejection of clusters that did not correspond to clean single units. Consistent with previous work ^20,33,40^, our results suggest that synthetic benchmarks should be designed to match the properties of the recording dataset and can be used to calibrate and test curation rules and thresholds against known ground truth.

Inferences about neural populations and network dynamics depend on both the quality and stability of the recorded signals and reliable identification of single units ^35^. Errors in single-unit identification can change individual spike trains as well as the composition and statistical structure of the recorded population. Over-splitting and duplicate detection can represent one neuron multiple times, inflating apparent population size and introducing artificial relationships or redundancy, whereas over-merging combines activity from distinct neurons and can distort estimates of neuronal tuning and temporal dynamics ^27^. Previous work showed that spike-sorting errors can produce biased estimates of firing rates and tuning that do not necessarily disappear with increasing amounts of data ^37^, and can systematically bias estimates of neuronal coincidence and correlation ^38^. This means that increasing the number of recorded neurons or behavioral trials does not necessarily correct systematic errors in unit identity. Our ground-truth simulations reveal an additional consequence of these errors at the population level. Duplicate and over-split clusters altered spike–spike relationships, while duplicate units introduced redundancy that reduced estimated population dimensionality (**Fig. 5**). These effects are particularly important for population analyses that depend on covariance structure across simultaneously recorded neurons to infer population organization and dynamics^36,48^. More data can reduce sampling variability, but it does not remove systematic errors in unit identity ^37,38^. Reliable single-unit identification should therefore be a prerequisite for investigating neural-circuit dynamics at both individual-neuron and population levels. We therefore propose treating curation as part of the recording workflow. Integrating curation with data acquisition can reveal problems in recording quality and inform improvements in the recording hardware and implantation protocol.

Several limitations remain. Synthetic recordings cannot reproduce the full biological and technical complexity of in vivo recordings ^33,40^. This is relevant for distinguishing related spike-sorting errors in neural data, such as over-splitting and duplicate re-detection, which cannot always be resolved unambiguously without ground truth. Hybrid ground-truth datasets can complement synthetic recordings by embedding known neuronal activity within real electrophysiological backgrounds ^20,31,34^. However, hybrid data provide ground truth only for the introduced units and cannot fully capture the interactions and complexity of the underlying biological population ^20,34^. UnitAssay also uses defined curation rules and thresholds that were developed for the recordings analyzed here. Whether the same thresholds generalize across other probe types, automated spike sorters other than Kilosort 4, and recording conditions remains to be tested. Within our dataset, however, UnitAssay was successfully applied across parietal, prefrontal, premotor, and motor cortices, demonstrating its use across distinct cortical regions. Our curation framework provides a way to recalibrate curation rules and thresholds using synthetic benchmarks matched to the recording dataset. MIND provides compact hardware for scaling multi-probe Neuropixels recordings while maintaining recording stability. However, the limits of scaling while preserving recording stability remain to be established. Future work should test stability over substantially longer durations, including week-long recordings, across larger simultaneously recorded neuronal populations and naturalistic, head-free recording conditions. Such experiments will establish how recording scale and duration can be increased while preserving recording stability and reliable single-unit identification. Future versions of MIND could also integrate Neuropixels recording with electrical or optical perturbation tools, enabling causal tests of neural-circuit dynamics. Together, MIND and UnitAssay provide a route toward scaling NHP electrophysiology while preserving recording stability and reliable single-unit identification.

## Methods

### Experimental preparation and behavioral task

We trained two adult male rhesus macaques (*Macaca mulatta*), Monkey G and Monkey T, to perform behavioral tasks involving eye- and hand-movements. Prior to behavioral training, each monkey underwent a surgical procedure to chronically implant a custom low-profile skull-chamber (Hybex Innovations Inc Canada) to facilitate head-fixation. The skull-chamber was designed to house a recording-chamber over the left hemisphere. Following completion of behavioral training, a second surgical procedure was performed to create the craniotomy and durotomy over the recording chamber for electrophysiological recordings. All experimental procedures conformed to NIH guidelines and were approved by the Institutional Animal Care and Use Committees (IACUC) at New York University and the University of Pennsylvania.

Each monkey performed a center-out reaching task. Each trial began with the appearance of a central target that the animal was required to hold for a variable delay (200–500 ms). A peripheral target was then presented in one of seven possible directions (0°, 45°, 90°, 135°, 180°, 225°, or 315° relative to the center). The monkey reached the peripheral target and maintained contact for 150–200 ms to receive a juice reward. Target size and acceptance windows were adjusted based on behavioral performance.

In addition, Monkey G performed a free-reach wand task, in which a handheld wand was presented at full reach distance by the experimenter. The monkey was trained to reach, grasp, and retract its arm without an explicit cue, receiving a juice reward upon successful reach. Arm position and movement trajectories were triangulated using synchronized camera recordings.

Monkey T also performed a two-target luminance-selection task. Each trial began with fixation on a central target for 500–800 ms, followed by extinction of the fixation point and the simultaneous presentation of two red peripheral targets (T1 and T2) located 10° from the center and separated by at least 90°. Target onset served as the go cue for a saccade toward one of the targets. Animal maintained fixation on the selected target for 300 ms to obtain a juice reward. Target luminance was randomized across trials, and in a subset of sessions, reward values for both targets were matched to ensure that target selection was primarily driven by exogenous visual factors rather than reward history.

### Multi-regional Implantable NeuroDrive (MIND)

All MIND components were custom-designed using Computer-Aided Design (CAD) software and fabricated in-house for bench testing. Components integrated into the implanted assembly were either machined (Protolabs) or 3D printed from biocompatible materials (3D SYSTEMS corporation or in-house using FormLabs 3D printer).

The Neuropixels hosting assembly consists of a compact microdrive capable of hosting one to six probes, which can be independently inserted. The microdrive incorporates a precise actuation mechanism, allowing for probe insertion in increments of 0.25 mm per leadscrew turn, within a compact 18 mm footprint. This system facilitates simultaneous multi-regional and multi-depth recordings. Actuation occurs when the leadscrew rotates through the threads of the shuttle, which is affixed to the dovetail of the Neuropixels probe (**Fig 1A**). Since the leadscrew head is fixed between the spacer and the screw guide, its vertical position remains constant during rotation. Consequently, the shuttle and connected probe move downward relative to the microdrive’s top surface (**Fig 1B, C**). The screw guide ensures stability by compressing the leadscrew head via a spring located between the screw head and the screw guide. Access to the leadscrew head is facilitated by a vertical hole in the screw guide, allowing insertion of a screwdriver without interference with the spring, thus enabling smooth actuation without significant jolting. The actuator block provides both protective housing and a sterilization compartment for the Neuropixels probes and shuttles, while also supporting the probes during brain tissue penetration. The bottom of the actuator block features a closed design, with small openings (approximately 1 mm in diameter) to accommodate the probe shanks.

The microdrive is available in various configurations, capable of hosting up to six Neuropixels probes, while keeping the microdrive height and width constant. The drive includes a notch in the actuator block, which allows for 360-degree rotation in 45-degree increments within the MIND insert (**Fig 1D**), enabling systematic mapping of accessible brain regions. The MIND insert integrates the microdrive into various MIND assemblies, each targeting different brain regions. A lock cap secures the microdrive within the insert, which is then connected to the electrophysiology extender. The electrophysiology extender is critical for accessing the targeted brain regions within different possible regions that are accessed in the chamber, as it enables the shifting of the opening window. The electrophysiology extender also adjusts the approach angle to the cortical surface, allowing approximately perpendicular probe penetration through region-specific extender designs. The electrophysiology extender designs are fixed onto the inner adapter, which securely anchors the artificial dura beneath. The artificial dura is attached beneath the natural dura at the chamber sides and held in place by the inner adapter. This configuration offers both protection to the brain tissue and stabilization of intracranial pressure, as the brain is enclosed within both the natural and artificial dura layers. The inner adapter is then affixed to the outer base, with the outer adapter part securely attached to the skull. The outer base provides access for cleaning the chamber, through suction ports, while the outer adapter allows for the attachment of a protective helmet to the skull, safeguarding the chamber and the entire assembly. This design enables electrophysiological recordings either with the animal’s head fixed (**Fig 1D**) or during free movement (**Fig 1E**). The head-stage platform in Fig. 1D ensures the stability of the Neuropixels cables during recordings, while having the cables supported by the connection to the datalogger for extended overnight recordings (**Fig 1E**).

MIND is compatible with both rodent and NHP Neuropixels probes and was used for implantation of both types. Because Neuropixels probes are fragile, especially during insertion, mechanical stability was a key consideration in our design. We observed that probes were more prone to breakage when inserted at acute angles relative to the cortical surface. To minimize mechanical stress and improve insertion success, probes were advanced perpendicularly. Probe tips were sharpened to ∼15° using a spinning hard-drive platter, consistent with previous reports ^9^. When necessary, the dura mater was peeled or thinned to facilitate smooth penetration.

Neuropixels 1.0 NHP probes, with their thicker shanks (90 µm), were more durable, whereas thinner rodent probes were more challenging to insert. To accommodate rodent probes, we incorporated a 2–3 mm agar gel layer into the actuator block, creating a gradual transition from agar to brain tissue and reducing bending during insertion. This modification was essential for successful implantation of rodent Neuropixels into the NHP cortex and demonstrates the cross-compatibility and adaptability of the MIND system across probe types and species.

### Data acquisition and collection

Neural signals were acquired using the SpikeGLX acquisition system at a 30 kHz sampling rate, with all probes internally time-stamped and synchronized within the same system. Behavioral task control was implemented through a custom Python-based graphical application that managed task events, reward delivery, and bidirectional communication with SpikeGLX to ensure precise temporal alignment. In a subset of experiments, eye position was monitored using a video-based pupil-tracking system, while arm and hand movements were captured using multiple synchronized high-speed cameras. A central task controller coordinated behavioral events, reward timing, and electrophysiological recording. Task events and synchronization pulses were digitally generated and routed through a National Instruments data-acquisition board, enabling alignment between behavioral, video, and neural datasets during offline analysis.

The dataset analyzed in this study comprised recordings collected across 20 experimental days, yielding a total of 147 Neuropixels recording sessions. A recording session corresponded to one SpikeGLX acquisition from one Neuropixels probe; during simultaneous multi-probe recordings, each probe contributed one recording session. Individual recording sessions lasted 2.0–36.5 min (median, 10.8 min). Of the 147 sessions, 26 were obtained from Monkey G and 121 from Monkey T.

Recordings were obtained using Neuropixels probes configured in either single-bank or multi-bank mode. In the single-bank configuration, recorded sites occupied a contiguous 3.82-mm span of the probe. In the multi-bank configuration, 383 recording sites were distributed across multiple banks, spanning 7.58–8.94 mm along the probe. Channels across multiple banks were selected using a classification-based selection (CBS) algorithm ^39^, allowing recording across an extended probe depth. Across the complete dataset, 75 recording sessions used a single-bank configuration and 72 used a multi-bank configuration.

Between one and four probes were deployed simultaneously using MIND. Nine experimental days used one probe, five days used two probes, five days used three probes, and one day used four probes, yielding 38 probe insertions across 20 experimental days. Recordings sampled posterior parietal cortex (PPC), prefrontal cortex (PFC), and premotor/motor cortex (PMd/M1), with 85 sessions from PPC, 14 from PFC, and 48 from PMd/M1. Overall, these recordings provided a dataset for evaluating MIND across single- and multi-probe recording conditions.

### Spike-sorting and curation pipeline

#### Preprocessing

Raw SpikeGLX binary files were preprocessed using CatGT (Janelia) before spike sorting. CatGT channel time-alignment correction was applied, followed by a 12th-order Butterworth band-pass filter from 300–9000 Hz (-apfilter=butter,12,300,9000). Global referencing was performed by subtracting the median signal across the 383 recording channels at each time point (-gblcar). Global referencing was used to preserve the spatially local signal structure subsequently used by UnitAssay for cluster-isolation analyses.

Transient artifacts were identified using CatGT’s *gfix* routine with thresholds of 0.40 mV for amplitude, 0.10 mV/sample for slope, and 0.02 mV for residual noise (-gfix=0.40,0.10,0.02). The routine identifies rapid excursions shared across a large fraction of recording channels. Flagged intervals were corrected independently on each channel by linear interpolation between the last unaffected sample before the artifact and the first unaffected sample after it. The thresholds were evaluated in a subset of recordings to confirm that they detected large, shared artifacts while remaining above the amplitude range of normal neural activity.

#### Kilosort 4 spike sorting

Automated spike sorting was performed using Kilosort 4 (v4.0.22). Each Neuropixels recording session was sorted independently using the same parameter configuration. Kilosort operated on the CatGT-preprocessed binary files using probe-specific channel maps reflecting the physical locations of the recorded contacts. Multi-bank recordings were sorted as a single 383-site probe, with discontinuities between recorded banks preserved in the channel map. All analyses reported in this study used the canonical dataset of 147 independently sorted recording sessions. Recordings from the same probe and insertion were additionally concatenated for illustrative analyses of recording stability, but concatenated sorts were not used for population statistics.

Kilosort was run with five drift-correction blocks (nblocks = 5), producing displacement estimates across nine overlapping depth blocks. The universal detection threshold (Th_universal) was set to 7.5 and the learned detection threshold (Th_learned) to 8.5. Th_universal controls spike detection during the initial pass using universal templates, whereas Th_learned controls spike detection during the final pass using templates learned from the data. Lower detection thresholds allow Kilosort to accept smaller events and can therefore alter the number of detected clusters, the number of spikes assigned to each cluster, and the composition of the resulting cluster population. Kilosort used 60,000-sample batches, a 5-µm drift-binning depth, and six principal components per channel. All other parameters were retained at their default values. Kilosort internal preprocessing, including a 300-Hz high-pass filter, common-average referencing, and whitening, was retained for all recordings. The stored Kilosort configuration was verified after sorting and was identical across all 147 recording sessions.

To evaluate the effect of detection thresholds on Kilosort output, we re-sorted 10 recording sessions of at least 20 min duration using five combinations of universal and learned detection thresholds. All other Kilosort parameters were held constant, and each sort was processed using the same UnitAssay configuration. UnitAssay was therefore used as a fixed downstream curation step to assess how changes in Kilosort detection parameters altered the resulting cluster population.

**Table 1:**
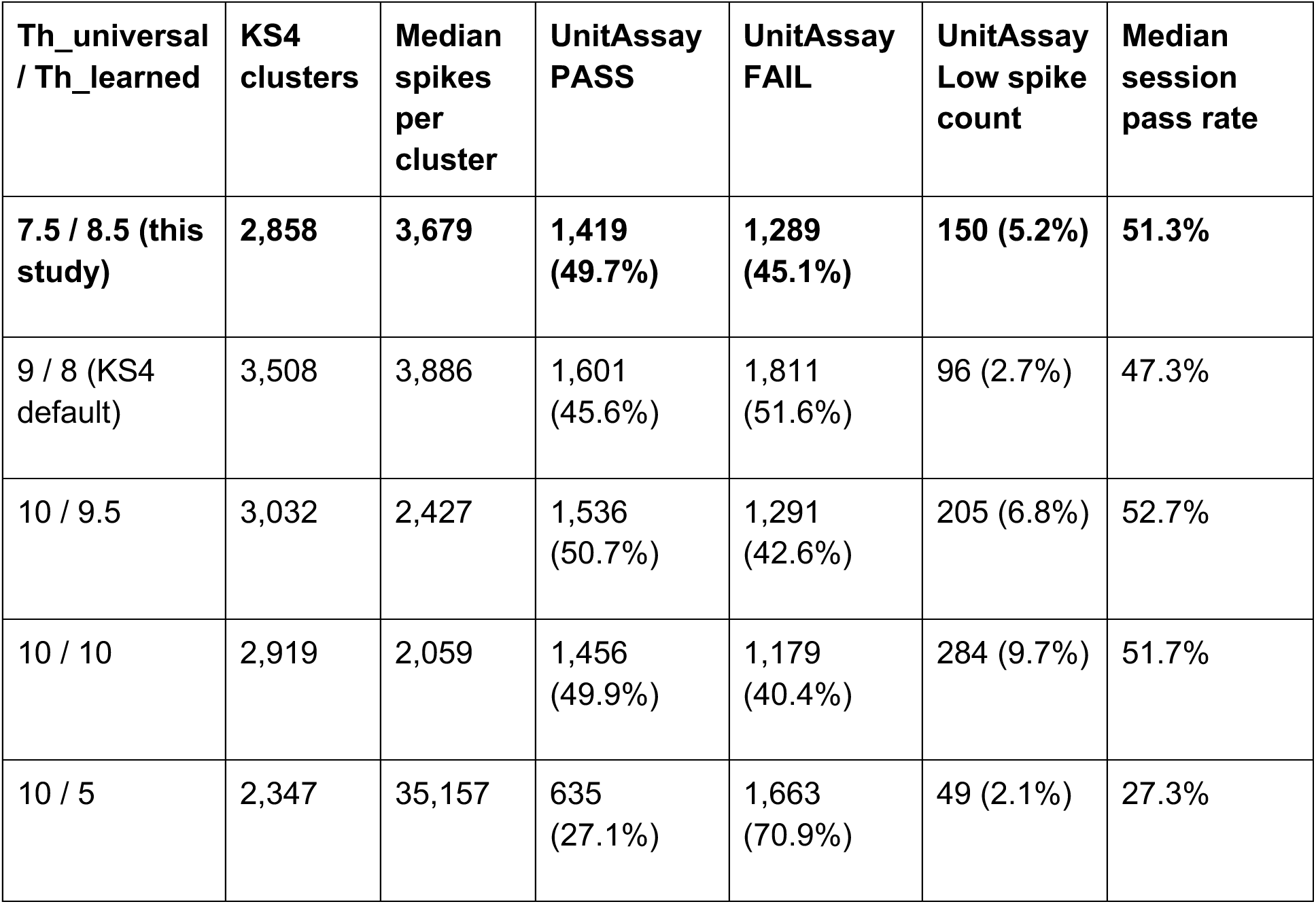
Effect of Kilosort 4 thresholds on cluster count.

| Th_universal / Th_learned | KS4 clusters | Median spikes per cluster | UnitAssay PASS | UnitAssay FAIL | UnitAssay Low spike count | Median session pass rate |
| --- | --- | --- | --- | --- | --- | --- |
| 7.5 / 8.5 (this study) | 2,858 | 3,679 | 1,419 (49.7%) | 1,289 (45.1%) | 150 (5.2%) | 51.3% |
| 9 / 8 (KS4 default) | 3,508 | 3,886 | 1,601 (45.6%) | 1,811 (51.6%) | 96 (2.7%) | 47.3% |
| 10 / 9.5 | 3,032 | 2,427 | 1,536 (50.7%) | 1,291 (42.6%) | 205 (6.8%) | 52.7% |
| 10 / 10 | 2,919 | 2,059 | 1,456 (49.9%) | 1,179 (40.4%) | 284 (9.7%) | 51.7% |
| 10 / 5 | 2,347 | 35,157 | 635 (27.1%) | 1,663 (70.9%) | 49 (2.1%) | 27.3% |

Across the 7.5/8.5, 9/8, 10/9.5, and 10/10 settings, UnitAssay retained approximately half of the clusters produced by Kilosort, with median session pass rates ranging from 47.3% to 52.7%. In contrast, the 10/5 setting substantially altered Kilosort output and reduced the median UnitAssay pass rate to 27.3%. These results emphasize that Kilosort detection thresholds should be selected carefully and kept consistent across recording sessions being compared.

#### Drift quantification

Probe displacement was estimated from the Kilosort 4 registration output (dshift), which reports vertical displacement over time for each spatial depth block. With nblocks = 5, Kilosort generates nine overlapping depth-block displacement traces. Drift analyses used the same 147 independently sorted recording sessions used throughout the study, comprising 75 single-bank and 72 multi-bank sessions. For each depth block, the displacement trace was centered on its median, and displacement was summarized as range₉₀, defined as the difference between the 95th and 5th percentiles of that block’s displacement trace. Thus, each recording session yielded nine block-specific range₉₀ values. A single session-level displacement value was obtained by taking the mean of range₉₀ across depth blocks. Group-level displacement values were summarized as the median of the session-level values across recording sessions. Displacement was compared with the 20 µm spacing between Neuropixels recording-site rows.

Single-bank and multi-bank configurations were defined from the spatial distribution of recorded contacts. Single-bank recordings occupied a contiguous 3.82-mm span of the probe, whereas multi-bank recordings spanned 7.58–8.94 mm and could contain unrecorded gaps between groups of contacts. Because Kilosort constructs its depth blocks from occupied depth indices without explicitly accounting for discontinuities in the channel map, some blocks in multi-bank recordings could fall within or span these unrecorded regions and contain relatively few recorded contacts. A block was classified as a gap block when its spatial extent contained fewer than 25% of the mean number of contacts per depth block. Single-bank recordings contained no gap blocks. Of the 72 multi-bank sessions, 51 contained at least one gap block. For the primary multi-bank analysis, gap blocks were excluded and the session-level displacement value was calculated as the mean range₉₀ across the remaining populated depth blocks. Displacement estimates calculated with gap blocks included were retained for comparison.

Sessions containing abrupt or coherent displacement events were distinguished from sessions showing continuous slow displacement. Artifact criteria were applied to sessions with range₉₀ greater than 15 µm. A session was classified as having a fast coherent displacement when an abrupt change greater than five times the median absolute inter-batch displacement occurred and involved, on average, at least 60% of depth blocks simultaneously. When fast coherent displacement occurred on two or more simultaneously implanted probes on the same experimental day, sessions from that day with range₉₀ greater than 15 µm were additionally flagged. Sessions not meeting these criteria were classified as having a step displacement when range₉₀ during the final 30% of the recording was ≤40% of that during the first 30%, indicating an early displacement followed by stabilization. Ten sessions met one or more artifact criteria, leaving 137 artifact-free sessions, including 68 single-bank and 69 multi-bank sessions.

Across the 68 artifact-free single-bank recording sessions, median range₉₀ was 4.0 µm (IQR, 2.6–7.0 µm), and all sessions remained below the 20 µm inter-row spacing. The relationship between probe displacement and recording duration was evaluated using Pearson correlation; displacement did not significantly increase with recording duration (*r* = 0.16, *p* = 0.20). For multi-bank recordings, displacement estimates were evaluated after excluding gap blocks, and recordings without gap blocks did not differ significantly from single-bank recordings (Mann– Whitney U test, *p* = 0.15).

#### Post-Kilosort processing

Kilosort output was converted into standardized data structures used for curation and downstream neural analyses. Spike times, cluster identities, template waveforms, principal-component features, spike-amplitude scaling factors, channel information, and Kilosort quality labels were extracted without modifying the original clustering assignments. For temporal analyses, cluster data were divided into consecutive non-overlapping 100-s windows spanning each recording. The same temporal grid was used for all clusters within a recording, and windows without sufficient spikes were retained rather than removed. Principal-component features from the target cluster, its local reference, and neighboring clusters were organized within these windows and subsequently used to quantify isolation stability. For behavioral analyses, spike times were aligned to task-relevant events and stored on a trial-by-trial basis. Reaching-task activity was aligned to movement onset, whereas eye-movement recordings were aligned to visual target onset. These data were used to generate peri-event rasters, firing-rate summaries, and population activity analyses.

### Automated curation: UnitAssay Layer 1 — screen

Layer 1 evaluates whether each Kilosort cluster is isolated from spatially local background activity and neighboring clusters, whether that isolation is maintained over the duration of the recording, and whether the cluster satisfies a refractory-period criterion.

#### Unit-channel and local-reference assignment

Each Kilosort cluster is associated with ten candidate channels on which Kilosort stores its principal-component features. For each candidate channel, waveform energy was calculated as the sum of squared template amplitudes, and spike representation (“dominance”) was calculated as the fraction of cluster spikes for which the first three principal-component features on that channel were not all zero. The unit channel was assigned as the highest-energy channel. If fewer than 50% of the cluster spikes were represented on that channel, the highest-energy channel with at least 50% dominance was selected instead. The local-reference channel was assigned as the lowest-energy channel with at least 10% dominance, non-zero waveform energy, and a waveform cosine similarity of ≤0.5 relative to the unit-channel waveform. If no candidate satisfied the waveform-similarity criterion, the lowest-energy channel satisfying the dominance and non-zero-energy requirements was used and the cluster was flagged. Channel assignment was performed independently for each cluster.

#### Feature representation

Layer 1 analyses were performed on a single channel at a time. Each spike was represented by its first three Kilosort principal-component features on that channel, multiplied by the spike-specific template scaling amplitude, producing a three-dimensional feature representation. For the local-reference comparison, the target cluster’s own spikes were represented using their features on the local-reference channel and compared with the same spikes represented on the unit channel. Neighbor comparisons included other clusters assigned to the same unit channel, with both clusters evaluated on that physical channel.

#### L-ratio

Cluster isolation was quantified using L-ratio ^41^. For each comparison population, the squared Mahalanobis distance of each comparison spike was calculated relative to the mean and covariance of the target-cluster distribution. L-ratio was defined as the summed upper-tail probability of these distances under a chi-square distribution with three degrees of freedom, normalized by the number of spikes in the target cluster:

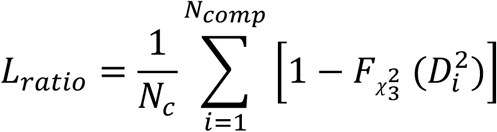

where 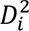 is the squared Mahalanobis distance of comparison spike *i* from the target-cluster distribution, 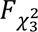 is the cumulative chi-square distribution with three degrees of freedom, *N*_comp_ is the number of comparison spikes, and *N_C_* is the number of spikes in the target cluster. Lower L-ratio values indicate greater separation. L-ratio was calculated independently between the target cluster and its local-reference population and between the target cluster and each neighboring cluster.

#### Isolation stability

To determine whether cluster isolation was maintained over time, L-ratio was recomputed independently in consecutive non-overlapping 100-s windows. A window was classified as isolated only when the local-reference comparison and all neighbor comparisons had L-ratio <0.2. Isolation stability was defined as the fraction of all windows satisfying this criterion. Windows in which the target cluster contained insufficient spikes to calculate its local-reference L-ratio were retained and counted as non-isolated rather than omitted. A cluster passed the isolation-stability criterion when at least 60% of all windows were isolated. Both the L-ratio threshold of 0.2 and the isolation-stability threshold of 60% are user-adjustable.

#### ISI-violation criterion

Refractory-period contamination was quantified using an observed-to-expected inter-spike-interval (ISI) violation ratio, following the refractory-violation framework of Hill et al., 2011 ^49^. An ISI shorter than 1 ms was considered a violation. For a cluster containing *n* spikes with firing rate *λ*, calculated over the interval between its first and last spike, the expected number of violations was

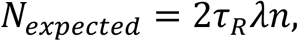

where *τ_R_* = 1 ms and *λ* = *n*/*T*, with *T* defined as the interval between the first and last spike of the cluster. The ISI-violation ratio was then calculated as

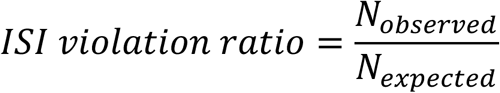

A cluster passed the refractory-period criterion when the violation ratio was <0.5 or when fewer than five violations were observed. Because genuine bursting neurons can produce short inter-spike intervals, clusters for which at least 80% of violations occurred at intervals <0.5 ms were classified as burst-like and passed the refractory-period criterion.

A cluster passed Layer 1 when it satisfied both the isolation-stability and refractory-period criteria.

### Automated curation: UnitAssay Layer 2 — curate

Layer 2 integrates multiple forms of evidence using a fixed sequential decision flow and assigns each cluster to the first rule that determines its outcome. The principal tests evaluate cluster composition, refractory-period violations, re-detection of the same spikes across clusters, contamination by local background activity, and separation from neighboring clusters. The rules were evaluated in the following order: composition → refractory → identity → clean → contamination → neighbor separation → stability. Composition was evaluated before identity because the re-detection test uses information about whether either member of a candidate pair is over-merged. Across 53,346 clusters from 147 recording sessions, the final UnitAssay outcomes were 22,482 PASS (42.1%), 24,493 FAIL (45.9%), and 6,371 LOW SPIKE COUNT (11.9%).

#### Feature-space separation

Layer 2 feature-space tests used the first three Kilosort principal components from each of the ten local channels, producing a 30-dimensional representation. Pairwise comparisons between clusters were restricted to channels shared by both clusters, with at least two shared channels required. Candidate groups were projected onto a regularized Fisher linear discriminant axis and tested for multimodality using Hartigan’s dip test ^50^. L-ratio was calculated in both directions, with each group’s spikes evaluated relative to the mean and covariance of the other group; the larger of the two L-ratios was retained. The number of degrees of freedom was three times the number of shared channels, and each group was required to contain at least 120 spikes to support covariance estimation. Two populations were considered separated when the dip-test significance was *p* < 0.05 and L-ratio was <0.05.

#### Composition

The composition test evaluates whether the spikes assigned to a single cluster contain evidence for more than one neuronal population using two complementary criteria: Hartigan’s dip test^50^ to test for multimodality and L-ratio^41^ to quantify separation between candidate populations in feature space. A cluster was classified as over-merged only when both criteria were satisfied (dip test, *p* < 0.05; L-ratio < 0.05).

A two-component Gaussian mixture model with diagonal covariance was first fitted to the cluster’s feature distribution to generate candidate subpopulations. If the smaller component contained fewer than four spikes, fired at <0.1 Hz, or contained fewer than 120 spikes, that component was set aside and the remaining population was evaluated. If the remainder contained fewer than 240 spikes, the composition test was not performed.

Candidate groups were projected onto a regularized Fisher linear discriminant axis and recursively subdivided when the dip test remained significant (*p* < 0.05), to a maximum recursion depth of three. Among the resulting groups containing at least 120 spikes, the pair with the lowest L-ratio was tested. Clusters satisfying both the dip-test criterion (*p* < 0.05) and the L-ratio criterion (<0.05) failed the composition test, except when the cluster had passed Layer 1 and the raw fraction of ISIs shorter than 1 ms was <0.0002. For these clusters, the candidate split was additionally tested for temporal ordering. A split was classified as time-ordered when the area under the ROC curve for spike time between the two feature-space groups exceeded 0.6, indicating a change in cluster composition over time. The composition test rejected 5,117 clusters because of two separable populations and an additional 179 clusters because of a time-ordered split.

#### Refractory

The refractory test3 used the same ISI-violation criterion defined in Layer 1. A cluster failed when the observed-to-expected ISI-violation ratio was ≥0.5, at least five violations were observed, and the cluster did not meet the burst criterion of ≥80% of violations occurring below 0.5 ms. This rule identified 5,818 clusters, of which 5,701 received a FAIL outcome and 117 were ultimately classified as LOW SPIKE COUNT.

#### Identity

The identity test evaluates whether two clusters represent re-detection of the same spike events. Clusters containing at least 30 spikes were compared with up to 25 nearby clusters sharing at least one footprint channel, defined as a channel on which both cluster templates exceeded 10% of their respective peak amplitudes. For each candidate pair, the modal temporal lag was identified over ±3 ms using 0.1-ms bins. Spike-train overlap was then calculated as the fraction of spikes in one cluster with a corresponding spike in the other cluster within ±0.5 ms of the pair-specific modal lag. A pair was classified as a re-detection when overlap exceeded the greater of a session-specific threshold or 50% and the absolute modal lag exceeded 0.2 ms. Thus, high overlap at or near zero lag alone was not sufficient to classify a pair as a duplicate.

The session-specific overlap threshold was derived from cluster pairs unlikely to represent the same neuron: their peak channels were separated by ≥400 µm and they shared no footprint channels. The threshold was set to the 98th percentile of this far-pair null distribution and was calculated separately for spike-count bands of 30–240, 240–700, 700–1,500, 1,500–3,000, 3,000–6,000, and >6,000 spikes. When fewer than 20 null pairs were available within a band, the pooled null distribution was used. For a re-detected pair, the cluster whose spikes were more completely contained within the other was rejected, unless only its partner was classified as over-merged. The identity rule identified 382 clusters; 215 received a FAIL outcome and 167 were ultimately classified as LOW SPIKE COUNT.

#### Clean

A cluster that passed Layer 1 and was not claimed by the composition, refractory, or identity tests was retained. This branch contained 20,561 clusters, of which 18,943 received a PASS outcome and 1,618 were ultimately classified as LOW SPIKE COUNT.

#### Contamination

Clusters that failed Layer 1 were routed according to whether the local-reference or neighbor comparison dominated the isolation failure. A cluster was routed to the contamination test when its local-reference L-ratio, averaged across time windows, was at least as large as the corresponding neighbor measure and reached 0.2 in at least one window. Because this evidence reflects overlap with the local background rather than with another identified cluster, no between-cluster comparison could clear the cluster. This rule contained 11,351 clusters, of which 9,974 received a FAIL outcome and 1,377 were ultimately classified as LOW SPIKE COUNT.

#### Neighbor separation

When Layer 1 failure was dominated by overlap with neighboring clusters, the target cluster was evaluated against every neighbor responsible for the isolation failure. An accusing neighbor was defined as a neighbor whose L-ratio averaged ≥0.2 across windows or reached ≥0.2 in at least 40% of windows. Feature-space separation was first evaluated using the dip-test ^50^ and L-ratio ^41^ criteria described above. A cluster was retained only when it could be separated from every accusing neighbor. This feature-space test retained 3,539 clusters and rejected 2,287 clusters; an additional 1,412 clusters reaching this branch were ultimately classified as LOW SPIKE COUNT.

Feature-space evidence was supplemented with independent time-domain tests. For each accusing pair, a cross-correlogram was calculated over ±25 ms using 0.5-ms bins. Expected coincidence was estimated from the mean count in flanking intervals 5–20 ms from zero, and deviations within ±2 ms were quantified relative to this baseline. The coincidence threshold was empirically calibrated using distant cluster pairs and varied with the spike count of the smaller cluster and the number of accusing neighbors. Threshold calibration covered smaller-cluster spike counts from 279 to 5,987 spikes and one to six accusing neighbors. Coincidence excess rejected 604 clusters, with an additional 153 below the final spike-count threshold.

A complementary refractory-deficit test asked whether the central cross-correlogram contained significantly fewer coincidences than expected from the flanking rate, consistent with one neuron having been split across two clusters. Significance was evaluated using a Poisson test, with a Šidák correction for multiple comparisons across neighboring clusters. This test rejected 304 clusters, with an additional 26 below the spike-count threshold. When one or more accusing neighbors could not be evaluated because of insufficient shared channels, insufficient spikes, or an uninformative cross-correlogram, the cluster was not cleared. Nine such clusters received a FAIL outcome, while 878 were below the final spike-count threshold.

#### Stability

Clusters that failed Layer 1 but had no window in which an L-ratio reached 0.2 and passed the refractory-period criterion were routed to the stability branch. These clusters failed because isolated windows constituted <60% of the recording, typically because the cluster was absent during substantial portions of the recording. This branch contained 726 clusters; 103 received a FAIL outcome and 623 were ultimately classified as LOW SPIKE COUNT.

#### Spike-count criterion

Clusters containing fewer than 240 spikes over the recording were reported as LOW SPIKE COUNT (not testable) regardless of the intermediate rule reached. The threshold of 240 spikes corresponds to twice the minimum of 120 spikes required for each population to support its own covariance estimate in the feature-space comparisons. Across the complete dataset, 6,371 of 53,346 clusters (11.9%) fell below this threshold.

#### Curation outcome and taxonomy

Curation decision and error classification were treated as separate operations. Layer 2 first assigned each cluster a PASS, FAIL, or LOW SPIKE COUNT outcome. A subsequent taxonomy integrated findings across the curation flow to assign an error class. PASS clusters were classified as clean. For failed clusters, findings were prioritized as low spike count, duplicate/re-detection, over-splitting, mixed/over-merged cluster, and intermittent presence. Because the error class incorporates findings across the complete flow rather than only the rule that issued the decision, counts for individual error classes need not equal the number of clusters rejected by the corresponding decision rule.

Across all 53,346 clusters, 22,482 (42.1%) were classified as clean, 22,678 (42.5%) as over-split, 6,371 (11.9%) as low-spike-count, 1,356 (2.5%) as mixed/over-merged, 357 (0.7%) as duplicates, and 102 (0.2%) as intermittently present.

#### Synthetic ground-truth data

UnitAssay performance was evaluated using 28 simulated extracellular recordings generated with MEArec through SpikeInterface ^28,40^, for which the identities and spike times of the underlying neurons were known. The simulations used Neuropixels 1.0 probe geometry and were designed as controlled scenarios that varied noise level, drift, bursting, unit density, firing rate, intermittent activity, refractory-period violations, and specific spike-sorting failure modes. Across the 28 recordings, 1,389 neurons were planted. Thirty-five placements corresponding to two poorly recovered templates were excluded from scoring, leaving 1,354 ground-truth neurons; 42 of these were not recovered by any Kilosort cluster. Kilosort 4 produced 5,018 clusters across the complete synthetic dataset. Of these, 4,758 could be assigned a ground-truth category, and 3,664 contained at least 240 spikes and were included in benchmark evaluation. All synthetic recordings were sorted using the same Kilosort 4 configuration and processed with the same UnitAssay pipeline used for the neural recordings.

#### Benchmark assessment

Kilosort clusters were matched to planted neurons using spike-train agreement, and ground-truth error categories were assigned from spike coincidences within a 1-ms window. Spikes assigned to a cluster but originating from another planted neuron were classified as over-merging, whereas unattributable spikes were classified as noise contamination; spikes from the same neuron appearing in additional clusters were classified as over-splitting. A contaminating neuron was required to account for at least 10% of a cluster’s spikes, and clusters with at least 50% unattributable spikes were classified as noise clusters. Of the 3,664 evaluable clusters, 1,134 were clean single units, 2,226 were over-split fragments, 127 were over-merged clusters, 155 were noise clusters, and 22 were noise-contaminated clusters. Benchmark evaluation was restricted to clusters containing at least 240 spikes. A true positive was a clean ground-truth single unit that was retained, a false positive was a non-clean cluster that was retained, a true negative was a non-clean cluster that was rejected, and a false negative was a clean single unit that was rejected. UnitAssay was compared with Kilosort 4’s Good/MUA labels and with a threshold-based quality-control procedure implemented in SpikeInterface (v0.104.8), applied to the identical set of clusters. SpikeInterface quality metrics included the ISI-violation ratio calculated using a 1.5-ms ISI threshold, presence ratio calculated in 60-s bins, and amplitude cutoff. A cluster passed the SpikeInterface comparator only when all three criteria were satisfied: ISI-violation ratio ≤0.5, presence ratio ≥0.8, and amplitude cutoff ≤0.1. Threshold boundaries were inclusive, and clusters for which any required metric could not be calculated (NaN) were classified as failures. Precision, sensitivity, specificity, F1 score, and balanced accuracy were calculated from the resulting confusion matrices.

#### Population correlations, dimensionality, and neural trajectories

To examine the effects of spike-sorting errors on population analyses, we calculated pairwise spike-count correlations using Pearson’s correlation coefficient (r) in non-overlapping 50-ms bins. Population dimensionality was estimated using the participation ratio (PR) of the eigenvalues (λ) of the correlation matrix, PR = (Σλ)²/Σλ², normalized by the number of units (n). Synthetic recordings were used to compare ground-truth neurons, Kilosort 4 output, curated units, and populations with duplicate or over-split errors selectively removed. Population correlation and dimensionality analyses used 25 of the 28 synthetic recordings, excluding three simulations that shared the same underlying recording. Duplicate clusters were identified when more than 50% of the smaller cluster’s matched spikes overlapped with those of another cluster assigned to the same ground-truth neuron.

For experimental recordings, curated single-unit activity (SUA) was compared with multi-unit activity (MUA) obtained from threshold crossings on individual Neuropixels channels. MUA was extracted from the same CatGT-preprocessed signals used for spike sorting, using a negative threshold of −4 times the channel-specific noise standard deviation (SD). Noise was estimated as the median SD across 60 one-second recording segments, with a 1-ms dead time between consecutive crossings. Channels with firing rates below 0.5 Hz and identified bad channels were excluded where applicable. Pairwise correlations were calculated using 50-ms bins, and PR/n was evaluated at matched population sizes (PMd: n = 261; PPC: n = 145), averaging across 50 random MUA channel subsets.

For neural trajectory analysis, spike counts were calculated in 10-ms bins and smoothed using a 20-ms Gaussian kernel. Activity was averaged across trials within each movement direction, normalized by the firing-rate range plus 5 Hz, and mean activity across directions was subtracted at each time point. Principal component analysis (PCA) was performed separately for SUA and MUA. The first three principal components were used to obtain neural trajectories, and MUA trajectories were aligned to SUA using orthogonal Procrustes transformation. Trajectories were visualized in the first two principal components around movement onset. Direction-specific structure was quantified using cross-validated R², predicting SUA trajectories from MUA trajectories with a leave-one-direction-out linear model across 20 random trial splits. The SUA reliability ceiling was estimated using independent trial halves from the curated population.

#### Computational resources

All processing was performed on a shared server containing two Intel Xeon Gold 6326 CPUs (2.90 GHz; 16 cores per socket, 64 total threads) and 755 GB RAM. Kilosort 4 was run using one NVIDIA RTX A4500 GPU with 20 GB memory per session, whereas CatGT and UnitAssay were CPU-only. Processing times were evaluated across 146 recording sessions with complete timing information for all pipeline stages. Median CatGT preprocessing time was 7.8 min per session (range, 1.1–22.5 min), and median Kilosort 4 processing time was 11.3 min (range, 1.5–49.7 min). UnitAssay required a median of 35 s for Layer 1 (range, 2–164 s) and 28 s for Layer 2 (range, 1–123 s), corresponding to a median total processing time of 62 s per recording session (range, 3–287 s). Median per-cluster processing times were 93 ms for Layer 1, 74 ms for Layer 2, and 166 ms for the complete UnitAssay pipeline. The median processing time for the complete CatGT–Kilosort 4–UnitAssay pipeline was 20.0 min per session, of which UnitAssay accounted for 4.9%. UnitAssay was run without GPU acceleration; Layer 1 was limited to eight CPU threads per recording session.

## Data availability

The data supporting the findings of this study are available from the corresponding authors upon reasonable request.

## Code availability

The UnitAssay code developed and used in this study is available at https://github.com/agritaD/UnitAssay

## Acknowledgements

We thank Marsela Rubiano for assistance with animal care and experimental procedures. This work was supported by grants R01-EY035826 and UF1-NS122123 to B.P.

## Author contributions

A.D. and B.P. conceptualized the project. A.D., J.C. and K.W. collected the experimental data. J.C., M.K., A.D., K.W., B.P. and J.S. contributed to the design and testing of MIND. A.D. and Q.X. developed UnitAssay and performed the associated analyses. A.D., J.H., and J.C. contributed software for behavioral and neural data acquisition and analysis. A.D. and Q.X. wrote the original manuscript. A.D. and B.P. reviewed and edited the manuscript.

## Supplementary figures

**Supplementary Figure 1.**
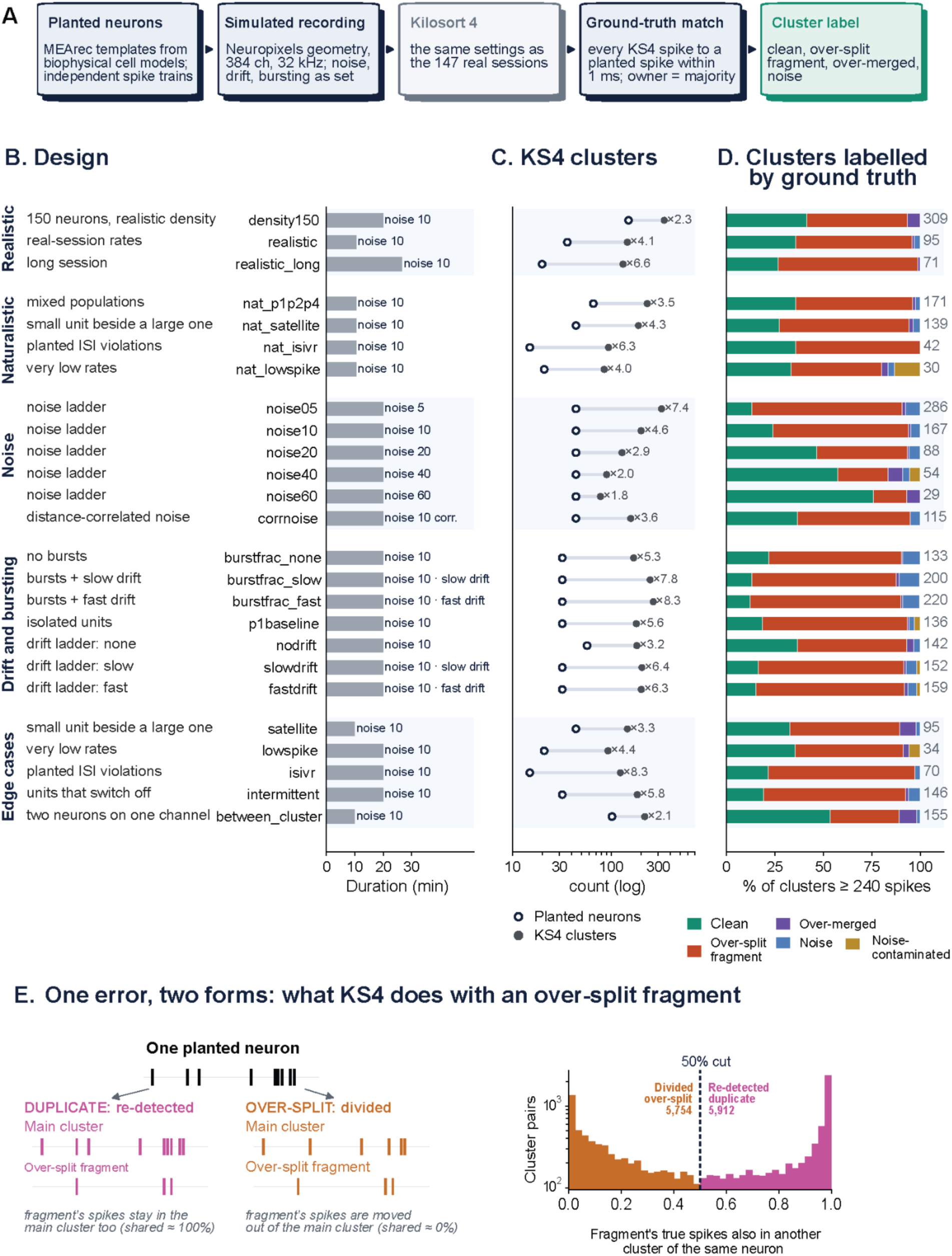
Design and ground-truth composition of the synthetic Neuropixels benchmark. **(A)** Synthetic-data generation and scoring pipeline. Independent spike trains from biophysically derived MEArec neuron templates were used to generate Neuropixels-like recordings with 384 channels and 32-kHz sampling. Noise, drift, bursting, and other recording properties were varied across simulations. Recordings were sorted with Kilosort 4 using the same settings as the neural recordings. Each Kilosort spike was matched to a planted neuron within 1 ms, and clusters were assigned a ground-truth label based on the matched spike identities. **(B)** Simulation design. Recordings spanned realistic, naturalistic, noise, drift-and-bursting, and targeted edge-case conditions. Bars indicate recording duration for each simulation condition; labels at right indicate the corresponding noise and drift settings. **(C)** Number of planted neurons and Kilosort 4 clusters for each simulation condition. Open circles indicate planted neurons and filled circles indicate detected Kilosort clusters. Values at right show the ratio of Kilosort clusters to planted neurons. **(D)** Ground-truth composition of Kilosort 4 clusters containing at least 240 spikes. Stacked bars show the fraction classified as clean, over-split fragments, over-merged, noise, or noise-contaminated; numbers at right indicate the number of evaluable clusters in each condition. **(E)** Two outcomes of a single over-split fragment. When most spikes in the fragment are also present in another cluster representing the same planted neuron, the fragment is interpreted as a re-detected duplicate; when the fragment contains complementary spikes that are absent from the main cluster, it is interpreted as a divided over-split fragment. Left, example spike trains illustrating the two cases. Right, distribution of the fraction of each fragment’s ground-truth spikes that are also present in another cluster from the same planted neuron. A 50% shared-spike criterion separates divided over-split fragments from re-detected duplicates.

**Supplementary Figure 2.**
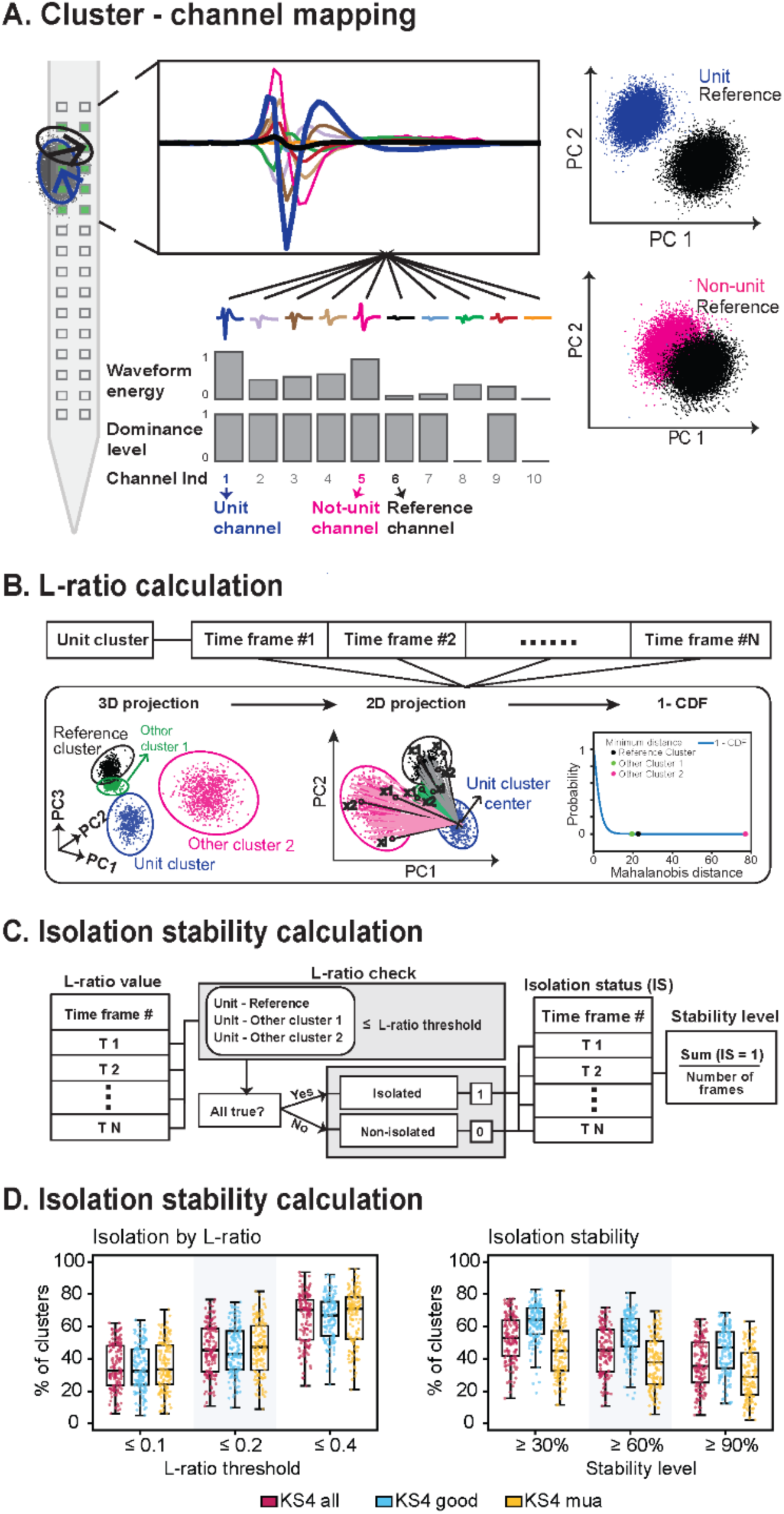
UnitAssay Layer 1 (screen). **(A)** Cluster–channel mapping. Each Kilosort 4 cluster is associated with ten candidate channels on the Neuropixels probe. Top: average spike waveforms across the ten channels. Bottom: waveform energy and spike-based dominance for each candidate channel. Waveform energy reflects the strength of the cluster waveform on each channel, while dominance reflects the representation of cluster spikes on that channel. A high-energy channel with sufficient dominance is assigned as the unit channel, while a lower-energy channel with sufficient cluster representation is selected as the local reference channel. Right: example principal-component feature spaces showing the unit cluster relative to its reference and a non-unit channel relative to the same reference. **(B)** L-ratio calculation. Cluster isolation is evaluated in principal-component feature space relative to the local reference and neighboring clusters. Spike distributions are projected from three-dimensional to two-dimensional feature space, and Mahalanobis distance is used to quantify separation. The complementary cumulative distribution (1−CDF) gives the probability that a spike at a given distance could arise from the unit-cluster distribution; lower L-ratio values indicate better isolation. **(C)** Isolation stability calculation. L-ratio is calculated independently in consecutive 100-s time windows. Within each time window, the unit cluster is compared with the local reference and neighboring clusters. A window is classified as isolated (IS = 1) only when all L-ratio comparisons fall below the defined threshold; otherwise, IS = 0. Isolation stability is calculated as the fraction of isolated windows across the recording. The default L-ratio threshold is 0.2, and the default isolation-stability criterion is 60%; both are user-adjustable. **(D)** Effect of isolation criteria on cluster yield across 147 recording sessions. Left: percentage of clusters meeting L-ratio thresholds of ≤0.1, ≤0.2, and ≤0.4. Right: percentage of clusters meeting isolation-stability criteria of ≥30%, ≥60%, and ≥90%. Results are shown for all Kilosort 4 clusters and separately for clusters labeled “Good” and “MUA.”

**Supplementary Figure 3.**
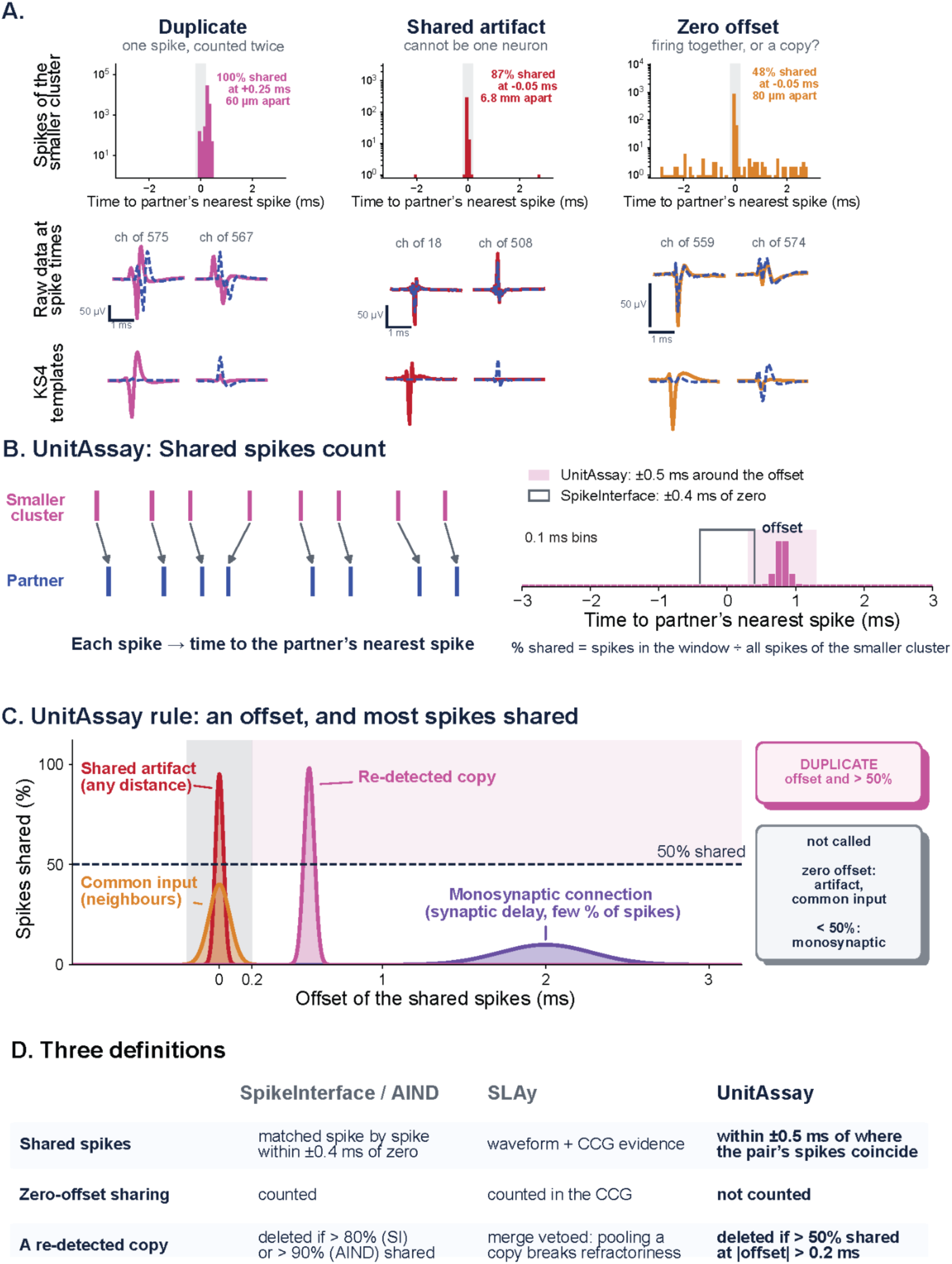
UnitAssay distinguishes re-detected duplicates from synchronous and shared-artifact spike trains. **(A)** Representative cluster pairs illustrating three forms of coincident spiking. Left, a re-detected duplicate in which the same spike events are assigned to two clusters with a consistent non-zero temporal offset. Middle, a shared artifact producing highly coincident activity across spatially distant channels. Right, a zero-offset pair in which coincident firing could reflect either a duplicated signal or genuine neuronal synchrony. Top row shows the distribution of each spike in the smaller cluster relative to its nearest spike in the partner cluster. Middle and bottom rows show raw voltage waveforms and corresponding Kilosort 4 templates at representative channels. **(B)** UnitAssay shared-spike calculation. For each spike in the smaller cluster, the nearest spike in the partner cluster is identified. The offset is estimated from the nearest-spike-time distribution, and shared spikes are counted within ±0.5 ms of that offset. Percent shared is calculated relative to the total number of spikes in the smaller cluster. The SpikeInterface comparison window around zero lag is shown for reference. **(C)** UnitAssay duplicate criterion. Cluster pairs are classified as re-detected duplicates when a consistent non-zero offset is present and more than 50% of spikes in the smaller cluster are shared with the partner. Zero-offset sharing is not classified as a duplicate because it can arise from shared artifacts, common input, or genuine synchrony. Pairs with fewer than 50% shared spikes are also not classified as duplicates, allowing short-latency physiological interactions such as monosynaptic coupling to remain unresolved rather than being removed. **(D)** Comparison of duplicate definitions used by SpikeInterface/AIND, SLAy, and UnitAssay. Rows summarize how each approach defines shared spikes, treats zero-offset sharing, and determines whether a cluster represents a re-detected copy.

**Supplementary Figure 4.**
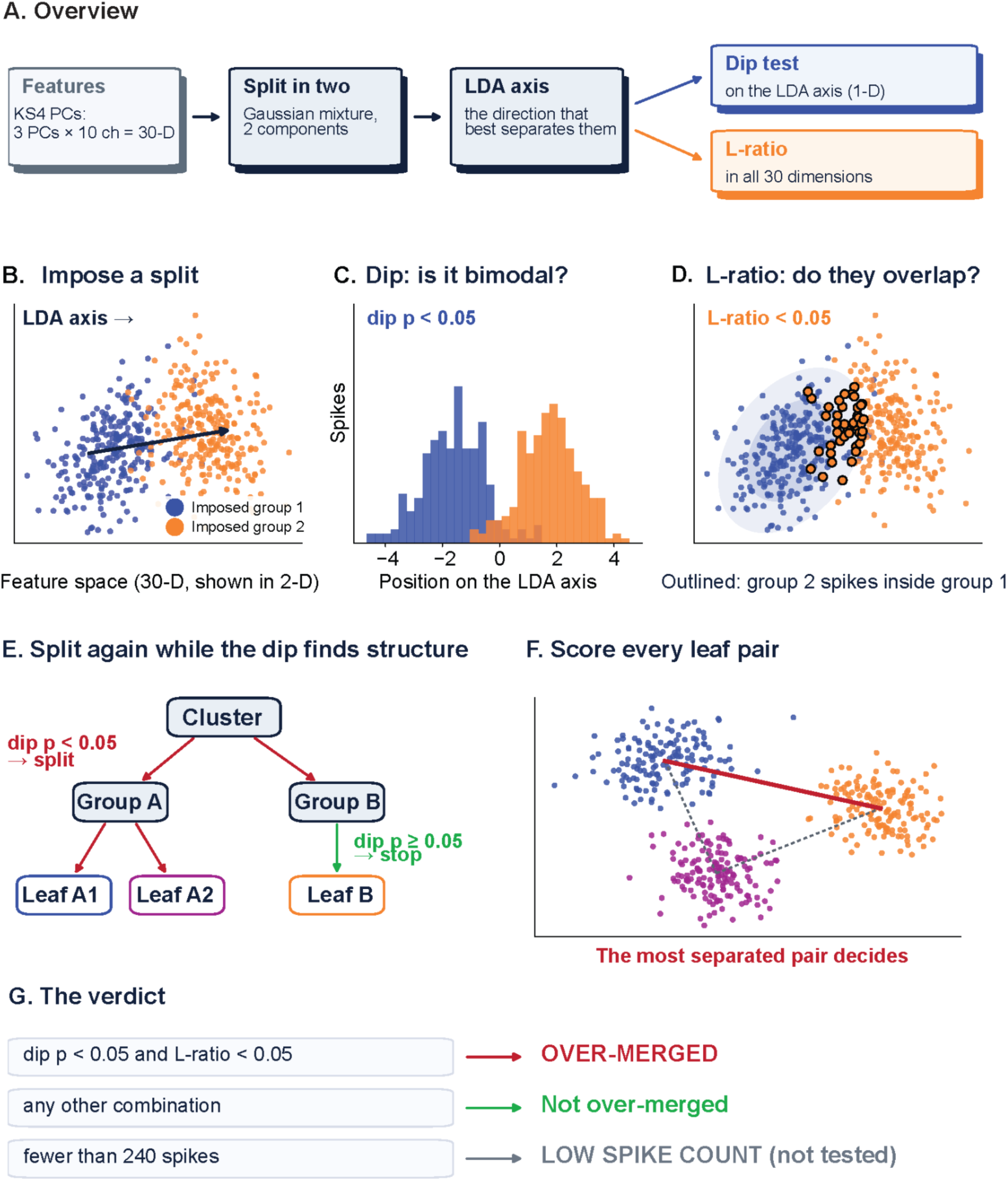
UnitAssay composition test for identifying over-merged clusters. **(A)** Overview of the composition test. Kilosort 4 principal-component features from three PCs across ten channels define a 30-dimensional feature space. Each candidate cluster is split into two groups using a two-component Gaussian mixture model. The linear discriminant analysis (LDA) axis that best separates the groups is then used for a dip test of bimodality, while L-ratio is calculated in the full 30-dimensional feature space to quantify overlap between the groups. **(B)** Example imposed split of a cluster in feature space. Points show the two groups defined by the Gaussian mixture model; the arrow indicates the LDA axis used to project the data. **(C)** Distribution of spikes along the LDA axis. Hartigan’s dip test evaluates whether the projected distribution is significantly bimodal; dip p < 0.05 indicates evidence for separable structure. **(D)** L-ratio evaluation of overlap between the imposed groups in the original feature space. Outlined points indicate spikes from one group that fall within the distribution of the other group. Lower L-ratio values indicate better separation; the example shown satisfies the L-ratio criterion of <0.05. **(E)** Recursive splitting procedure. When the dip test identifies significant structure, the corresponding group is split again; splitting stops when no further bimodality is detected. This produces a set of terminal leaf groups. **(F)** Pairwise comparison of all terminal leaf groups. Separation is scored for each leaf pair, and the most strongly separated pair is used to determine the composition outcome. **(G)** Final decision rule. A cluster is classified as over-merged when the most separated leaf pair satisfies both dip p < 0.05 and L-ratio < 0.05. Clusters not meeting both criteria are classified as not over-merged. Clusters with fewer than 240 spikes are not tested and are labeled low spike count.

